# Confidence is predicted by pre- and post-choice decision signal dynamics

**DOI:** 10.1101/2023.01.19.524702

**Authors:** John P. Grogan, Wouter Rys, Simon P. Kelly, Redmond G. O’Connell

## Abstract

It is well established that one’s confidence in a choice can be influenced by new evidence encountered after commitment has been reached, but the processes through which post-choice evidence is sampled remain unclear. To investigate this, we traced the pre- and post-choice dynamics of electrophysiological signatures of evidence accumulation (Centro-parietal Positivity, CPP) and motor preparation (mu/beta band) to determine their sensitivity to participants’ confidence in their perceptual discriminations. Pre-choice CPP amplitudes scaled with confidence both when confidence was reported simultaneously with choice, or when reported 1-second after the initial direction decision. When additional evidence was presented during the post-choice delay period, the CPP continued to evolve after the initial choice, with a more prolonged build-up on trials with lower confidence in the alternative that was finally endorsed, irrespective of whether this entailed a change-of-mind. Further investigation established that this pattern was accompanied by earlier post-choice CPP peak latency, earlier lateralisation of motor preparation signals toward the ultimately chosen response, and faster confidence reports when participants indicated high certainty that they had made a correct or incorrect initial choice. These observations are consistent with confidence-dependent stopping theories according to which post-choice evidence accumulation ceases when a criterion level of confidence in a choice alternative has been reached. Our findings have implications for current models of choice confidence, and predictions they may make about EEG signatures.

## Introduction

Mathematical modelling and neurophysiological investigations of perceptual decision making suggest that choice confidence evolves over the course of deliberation, and is informed by several factors, including the strength of evidence favouring the chosen alternative (Bang & Fleming, 2018), the difference in evidence between the available options (Li & Ma, 2020), and the time taken to decide (Kiani et al., 2014). Confidence judgments can also be updated after a decision has been made, taking account of evidence accumulated while the decision-reporting action is still being executed (Resulaj et al., 2009) and/or new evidence encountered after response completion (Fleming et al., 2018; Moran et al., 2015; van den Berg et al., 2016; Yu et al., 2015). What form such post-commitment accumulation processes take remains unclear and several alternative possibilities have been proposed.

According to some models, confidence levels are updated by a continuation of the same evidence accumulation process that informed the initial choice (Moran et al., 2015; Pleskac & Busemeyer, 2011; Yu et al., 2015) but other accounts invoke a distinct metacognitive process that evaluates the accuracy of the preceding choice (Desender, Ridderinkhof, et al., 2021; Fleming & Daw, 2016). Another key point of distinction between existing models of confidence in perceptual decisions, is whether they assume that post-choice evidence accumulation operates until confidence is probed (Yu et al., 2015), or is terminated when a criterion level of confidence (Pleskac & Busemeyer, 2011) or elapsed time (Desender, Donner, et al., 2021) has been reached (‘optional stopping’). Adjudicating among these accounts has been difficult as post-choice evidence accumulation has rarely been observed or probed in the brain with sufficient temporal precision. Electrophysiological research in monkeys using opt-out paradigms has established that choice confidence can be read-out jointly from the pre-choice firing rate of decision variable encoding neurons and deliberation time (Kiani & Shadlen, 2009), but this work has not yet examined the neural underpinnings of post-choice confidence reports.

One promising signal for probing post-choice confidence representations in humans is the Centro-parietal positivity (CPP), which tracks sensory evidence accumulation in the lead up to a decision (O’Connell & Kelly, 2021). Whereas effector-selective decision signals previously reported in humans and other species reach a stereotyped amplitude immediately prior to response execution, the CPP’s amplitude varies with several factors known to influence choice confidence, for example having greater amplitudes on correct trials and reduced amplitudes for trials with longer RTs (Kelly et al., 2021; Steinemann et al., 2018). Thus far, only a few studies have directly investigated the CPP’s relationship with confidence and these have yielded apparently mixed results. While several studies have reported positive relationships between pre-choice CPP amplitude and confidence (Davidson et al., 2021; Gherman & Philiastides, 2015, 2018; Herding et al., 2019; Pereira et al., 2021; Tagliabue et al., 2019), two studies found no such relationship (Feuerriegel et al., 2022; Rausch et al., 2020).

Elsewhere, it has been reported that a post-choice Centro-parietal signal with identical topography to the CPP, but traditionally labelled as the Error Positivity (Pe), also scales with choice confidence but in the opposite direction to that reported for the pre-choice CPP. The Pe is seen after erroneous choices that are explicitly reported to be incorrect (Falkenstein, 1990; Nieuwenhuis et al., 2001; Steinhauser & Yeung, 2010), and its amplitude increases the more certain participants are that they have made an error (Boldt & Yeung, 2015; Feuerriegel et al., 2022). The Pe also exhibits a build-to-threshold relationship with error signalling reports (Murphy et al., 2015), and predicts subsequent post-error slowing and post-choice information-seeking (Desender, Boldt, et al., 2019; Desender, Murphy, et al., 2019).

While the Pe literature seems to suggest an “error accumulation” process gathering evidence for reversing a choice, much of this research involved studies with no post-choice evidence available (Boldt & Yeung, 2015; Desender, Donner, et al., 2021) or where participants only responded when errors were detected (Murphy et al., 2015). To our knowledge, there has not been a direct comparison of the post-commitment dynamics of neural evidence accumulation signals with versus without continued evidence presentation. Additionally, recent work has highlighted that the use of baseline correction to a time before just before the initial response may have caused pre-choice amplitude differences to be transferred to post-choice amplitude measurements (Feuerriegel et al., 2022). Finally, this previous work has not considered the important influence that optional stopping could have on trial-averaged signal amplitude measurements and their interpretation. Here, we first demonstrate that when dot motion direction decisions and the participant’s associated confidence are reported simultaneously, pre-choice CPP amplitude scales with confidence. In a second experiment, we used a contrast discrimination task in which participants withheld their confidence reports for a one-second post-commitment delay, and randomly varied across trials whether the physical evidence remained on screen or was extinguished. This manipulation allows us to demonstrate that despite physical evidence remaining available, post-choice accumulation of this evidence terminates early when participants achieve high confidence that they have identified the correct choice alternative, irrespective of whether or not this entails a reversal of their initial choice.

## Methods

### Ethics

The study was approved by Trinity College Dublin ethics committee and carried out in accordance with the Declaration of Helsinki, and EU GDPR. Written informed consent was given before the start of the first session.

### Participants

Participants were between 18 and 32 years old, with normal or corrected-to-normal vision, no history of neurological or psychiatric disorders, epilepsy, or unexplained fainting. Experiment 1 recruited 27 participants (13 female, 14 male), with two excluded following artefact rejection (see below). Experiment 2 recruited a different group of 30 participants (18 female, 12 male), five of whom were excluded due to insufficient data retained after artefact rejection. Participants were paid for their time (€35 in experiment 1, €45 in experiment 2), plus a bonus depending on their performance (experiment 1: up to €6.50, mean = €4.40, SD = 1.10; experiment 2: up to €16, mean = €8.50, SD = 1.50).

### Experimental design

#### Experiment 1 – Simultaneous Choice and Confidence Reports

Experiment 1 was a random dot-motion direction discrimination task in which participants simultaneously reported the direction of coherent motion and their choice confidence (Figure 1). The task was programmed in MATLAB (R2013b) and Psychtoolbox-3 (Kleiner et al., 2007). Testing took place in a dark, sound-attenuated room, with participants seated 57cm from a CRT monitor (51cm, 65.2cd/m^2^ luminance, 75 Hz refresh rate, 1024×768 resolution), with their head on a chin rest.

**Figure 1.**
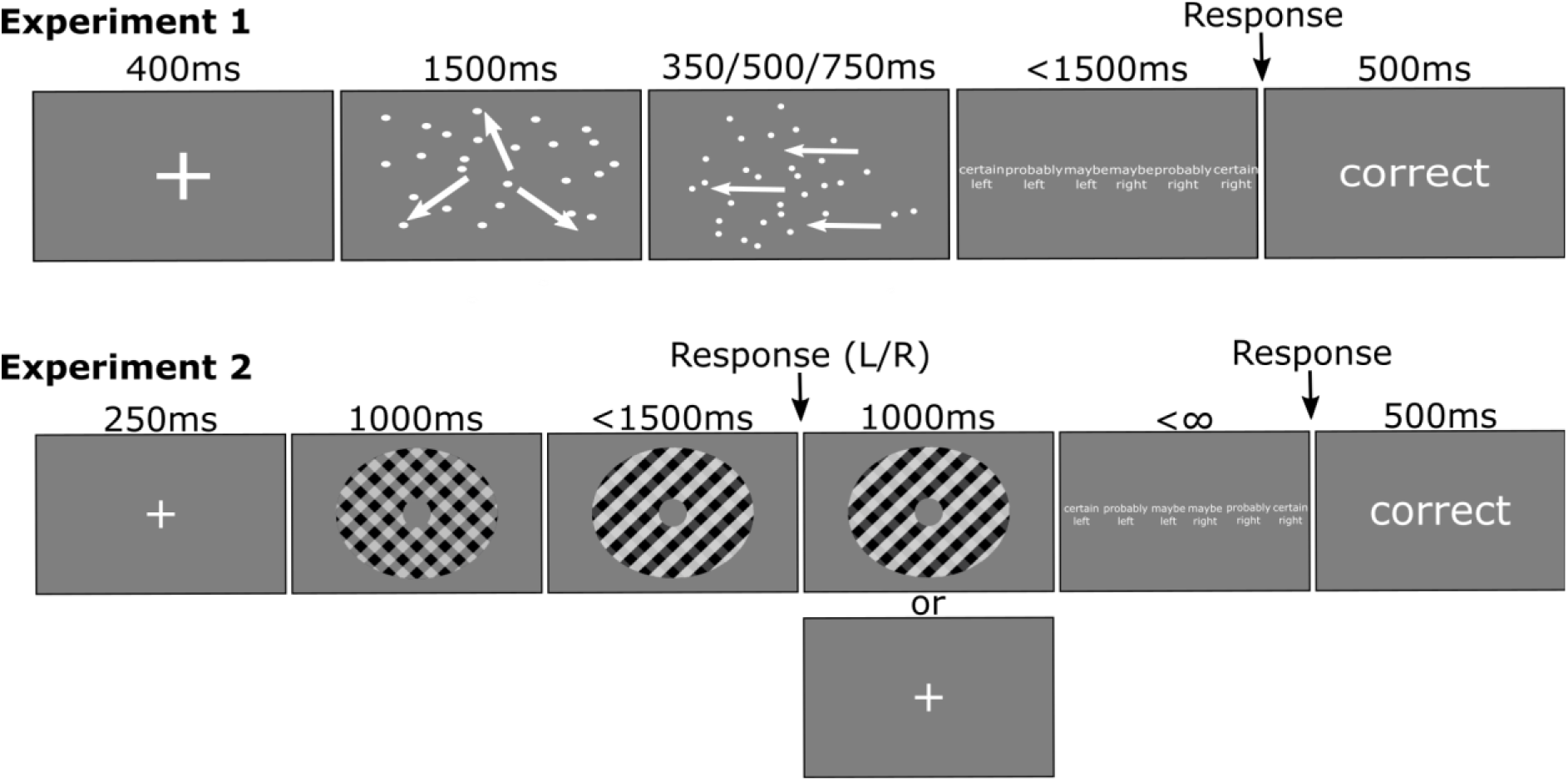
Task designs for experiments 1 and 2. Experiment 1 was a dot motion direction discrimination task, with simultaneous direction and confidence reporting. Each trial started with an initial 1500ms lead-in of incoherent motion to allow visual-evoked potentials associated with stimulus-onset to resolve, offering a clear view on choice-related signals. Participants withheld reporting their choice and confidence until the appearance of a 6-point confidence scale following stimulus offset (350/500/750ms stimulus duration). In experiment 2, participants were presented with overlaid gratings and reported whether the left or right grating was higher in contrast. Again, trials started with an initial zero-evidence lead-in period during which the gratings appeared at equal contrast. Initial left/right choices had to be reported within a 1500ms deadline. Participants were then probed to report their choice confidence using the same scale as in Experiment 1 after a post-choice delay of 1000ms. During the delay period the evidence either remained on screen (as shown in the figure) or was extinguished with equal probability. Note that stimulus sizes in all panels are not to scale

Trial onsets were self-paced, beginning when participants pressed the space button. A white central fixation cross was presented for 400ms, followed by the random dot kinematogram composed of 75 white dots (0.16° diameter) in an aperture (8° diameter) on a grey background. The dots moved with 0 coherence for an initial lead-in period of 1500ms to prevent visual-evoked potentials elicited by stimulus onset from overlapping with choice-related signals. A proportion of the dots began moving coherently after this 1500ms lead-in with the proportion individually titrated to achieve a criterion discrimination accuracy level (see below). Dot positions were updated on every frame, with a proportion (matching the coherence) randomly selected to move either left or right on each trial (equal probability) by 0.2° relative to their location three frames earlier, to give an overall motion speed of 15°s^-1^. The remaining dots were moved to a random new location every frame, and the coherent dots were re-selected every frame to prevent people tracking individual dots. The coherent motion was presented for 350ms, 500ms, or 750ms (equal probability), and was followed by the appearance of a 6-point response scale. Participants rested the first three fingers of each hand on the response keys of a keyboard and were instructed to click the “s”, “d or “f” key with their left hand to indicate “certain left”, “probably left” or “maybe left” respectively and clicking “j”, “k”, “l” with their right hand to report “maybe right”, “probably right”, “certain right”, respectively. Participants were instructed to withhold reporting their choices until the appearance of the confidence scale and only responses within 0-1500ms of scale onset were recorded. Visual feedback was then provided for 500ms in the form of “correct”, “error”, “too fast” (RT<0ms) or “too slow” (RT>1500ms). Participants were shown their mean accuracy and response time at the end of each block and were informed of their bonus winnings at the end of the experiment (see scoring rule in Experiment 2 section).

Participants completed two consecutive days of testing, with training on the behavioural task and task difficulty titration taking place on day 1. Training started with 50 trials at high coherence, until participants performed close to 100% accuracy on the direction decision and were comfortable rating their confidence simultaneously. The coherence was then titrated in blocks of 30 trials to achieve approximately 75% discrimination accuracy using a staircase procedure that increased the coherence 1% after every error and decreased it by 1% after three consecutive correct responses (average titrated coherence 11.19%, SD = 4.66). The EEG testing session took place on Day 2 and consisted of 8 blocks of 72 trials with a short rest break in between blocks. Only the data from Day 2 are included in the analyses reported below.

#### Experiment 2 – Delayed Confidence Reports With or Without Post-Choice Evidence

Experiment 2 (Figure 1) was a contrast discrimination task in which participants made an initial speeded two-alternative choice and were subsequently cued to report their final choice confidence after a 1 second delay. In randomly interleaved trials, evidence was either Extinguished immediately after the initial choice report or Continued throughout the delay period. The same physical set-up was used as for experiment 1, except the monitor was a 40.5cm CRT monitor.

After pressing the space bar to begin the trial, participants saw a central fixation for 250ms, followed by two overlaid gratings tilted at 45° from vertical in each direction (spatial frequency = 2 cycles per degree) in an annulus shape around a central fixation (inner radius = 1°, outer radius = 6°). The two gratings were initially presented at 50% contrast. Evidence onset occurred after 1000ms, with one grating increasing by a criterion amount (see below) and the other grating decreasing by the same amount. To allow reverse correlation analyses (analyses not reported here) we introduced small frame-to-frame variations in the contrast-difference between the two stimuli with values drawn from a Gaussian noise distribution (SD=0.015 which corresponds to around 10% of the mean evidence strength, maximum variation was capped at 3 SD). Throughout the stimulus presentation, the two gratings were flickered at 15 Hz in anti-phase, which allowed us to recover a 15 Hz steady-state visual evoked potential (SSVEP) driven by the contrast difference between alternating gratings, whose phase indicated the direction of the difference. In addition, activity at 30 Hz indexed the summed visual response to the two gratings, offering a general index of engagement. Participants indicated which grating had the higher contrast with a button press (“f” or “j” for left or right) as soon as they liked, with a deadline of 1500ms. Following initial response, the evidence either Continued at the same mean strength or was Extinguished and replaced with a fixation cross. These trial types were randomly interleaved and occurred with equal probability. 1000ms after the initial response, the confidence rating scale was shown on the screen, and participants responded as in experiment 1 – importantly, the scale still ran from “certain left” to “certain right”. Participants were instructed to report their final choice confidence in light of all the evidence they had viewed, as opposed to retrospectively reporting on their confidence in their initial choice. Feedback was given as in experiment 1.

Participants completed 16 blocks of 80 trials while undergoing EEG recordings with half of the trials completed on the first day of testing, and half on the second (consecutive) day. These two days of data were combined for the analyses. The first day also included training and staircasing, using a two-down one-up staircase to reach 70% accuracy (step-size starting at 6% and decreasing by 1% point each step until they reached 1%; mean contrast = 13.19%, SD = 4.61). A bonus was paid out based on their accuracy and confidence in the main task, based on a quadratic scoring rule (points = 100*(1 – (initial accuracy – confidence)^2^)), scaled to €0-€16. Initial accuracy in this equation is coded as 1 for correct and 0 for incorrect, and confidence is expressed relative to initial choice on each trial, taking one of 6 equally spaced values ranging from 0, meaning ‘certain’ in the initially unchosen option, to 1, meaning ‘certain’ in the initially chosen option. This scoring rule incentivises both initial accuracy and the accuracy of confidence responses (Staël von Holstein, 1970) and also orthogonalizes confidence from expected reward; more points are given for high-confident accurate responses and low-confidence errors, and the maximising strategy is to have high initial accuracy and high metacognitive accuracy. We checked they understood this rule by asking them how many points they would get for a confident error.

### EEG acquisition and pre-processing

EEG was recorded with a BioSemi ActiveTwo system (BioSemi, Netherlands), with 128 scalp electrodes at 512 Hz. Vertical EOG was recorded from electrodes above and below the left eye. Data were processed and analysed using custom MATLAB scripts that drew on routines from EEGlab (Delorme & Makeig, 2004). EEG data were detrended, low-pass filtered at 40 Hz (FIR filter), epoched in intervals of −1000:3000ms relative to evidence onset, and baseline-corrected relative to an interval immediately preceding evidence onset (200ms in experiment 1; 400ms in experiment 2). The same pre-evidence baseline was used for pre- and post-choice CPP analyses in Experiment 2, except for the pre-response baseline investigation which used the 100ms before initial RT. Channels with extreme variance or high artefact counts were interpolated.

Epochs with scalp electrode voltages over 100μV, or with bipolar VEOG voltages over 200μV (250μV in experiment 2) between the baseline and the response onset (confidence cue onset in experiment 2) were flagged as artefactual and removed. Participants with more than half their trials removed were excluded from the analysis (2 in experiment 1, 5 in experiment 2). Response-aligned epochs were also extracted from the evidence-aligned epochs, using the interval −1000ms:500ms relative to response in experiment 1, and −1000ms:1000ms in experiment 2. The voltages were transformed into Current Source Density (CSD) using the CSDToolbox (Kayser & Tenke, 2006).

### Analysis

For both experiments, trials with initial choice reaction times less than 100ms were excluded. Because in experiment 2 participants were able to change their minds between the initial choice and the final confidence report, we scored their final reports in two different ways. First, to test the hypothesis that the post-choice CPP reflects a distinct accumulation process selectively gathering evidence calling for a revision of the initial choice, the final confidence ratings were recoded relative to the initial choice (herein referred to as ‘confidence-in-initial-choice’) taking into account whether there was a change-of-mind (CoM) or not. For example, an initial ‘left’ response followed by a ‘certain right’ was labelled ‘certain CoM’, while an initial left response followed by a ‘probably left’ was labelled as ‘probably no-CoM’. Note that, although we re-coded the confidence ratings relative to the initial choice, participants were instructed to rate their *current* confidence at the end of the trial as opposed to retrospectively reporting their confidence at the time of the initial choice. Secondly, we sought to test the hypothesis that the CPP scaled with confidence in the final choice report, irrespective of whether it conflicted with the initial choice or not (herein referred to as ‘confidence-in-final-choice’). Thus, trials on which participants reported ‘certain left’ or ‘certain right’ would be pooled together and labelled as ‘certain’.

Data were analysed with trial-wise linear mixed models (LMM), after z-scoring all variables, and including a random intercept of participant. The regression coefficients from these models are standardised by the z-scoring, and the degrees of freedom represent the total number of trials minus the degrees taken up by the factors included in the model. Generalised LMM were used for logistic regression when binary variables such as accuracy or change-of-mind were the dependent variable. Reaction times (RTs) were measured from stimulus offset in Experiment 1 (to account for different stimulus durations), and from evidence onset in Experiment 2. Experiment 2 final RTs were measured from the confidence-scale onset. Experiment 1 RTs and Experiment 2 final RTs were log-transformed for statistical analysis (Experiment 2 initial RTs were approximately normal). For experiment 1, Stimulus Duration was included in the LMM. The main factors of interest were Initial Accuracy and Confidence-in-final-choice, and experiment 2 also had Post-choice Evidence Condition (Continued or Extinguished) and Change-of-mind.

‘Maybe’ responses were less common than the others, especially when evidence was extinguished in Experiment 2, so a minimum number of 10 trials for each Confidence*Evidence combination was applied. This led to the exclusion of ‘maybe’ extinguished-evidence trials from 5 participants (35 trials in total; their other trials were kept in). Similarly, 7 participants in Experiment 1 had fewer than 10 trials for some Confidence*Stimulus Duration combinations, so these trials were excluded (46 in total). Other trials from these participants were kept in the analysis, as the LMM allows for missing data – the excluding these participants entirely did not change the pattern of results.

The added stochastic contrast variation in Experiment 2 intended for reverse correlation analyses was unable to replicate previous effects of initial choice accuracy, indicating that the variance was too low, and so also had no detectable effects of confidence ratings, so is not included here.

### EEG Signals

To identify appropriate electrodes for measuring the CPP, we examined the grand-average ERP topographies including all trials and covering the time range from −150:-50ms before the initial choice report. A cluster of 5 Centro-parietal channels centred on the focus of the CPP topography were selected and, for each individual, we identified the channel within this cluster that exhibited the largest pre-response amplitude in order to extract CPP measurements. The pre-choice mean amplitude was taken within −150:-50ms before initial response in Experiment 1, and −140:-50ms for Experiment 2 (in order to capture an integer number of cycles of the SSVEP signal). Examination of the post-choice CPP waveforms indicated that this signal dropped partially toward baseline soon after the initial choice report before undergoing a second build-up in advance of the presentation of the confidence cue, which differed between Continued and Extinguished post-choice evidence conditions towards the end of the delay (see Supplementary Figure 1). Based on this observation, we measured the amplitude of the post-choice CPP in the window 700:1000ms after the initial choice. A 10 Hz low-pass filter is applied for plotting only, to remove the 15 Hz and 30 Hz SSVEP components. We additionally examined the effect of a pre-response baseline (−100:0ms before initial response) on the post-choice CPP both in our time-window, and an earlier window (200:350ms) as reported in a previous paper (Feuerriegel et al., 2022). CPP post-choice peak latency was calculated on data low-pass filtered at 6Hz, by finding the latency of the maximal amplitude from 0:1000ms after initial response.

Effector-selective motor preparation was analysed via mu-beta band (8-30 Hz) activity. Fast-Fourier transforms were performed on the CSD data, with 256-sample windows (~500ms) moving in 8-sample steps (20ms). Clusters of three electrodes in each hemisphere were chosen, centred on the strongest ipsilateral minus contralateral pre-response (−150:-50ms) amplitude in the grand-average topography. Within these clusters, the channel with the largest difference in amplitude was chosen for each person. Mean amplitudes were calculated for the interval −250:-100ms before initial choice in experiment 1, and −300:0ms in experiment 2, along with a post-choice measurement in experiment 2 from 700:1000ms (i.e., −300:0ms before confidence cue appears).

In experiment 2, the two gratings each flickered at 15 Hz in anti-phase generating distinct phase-tagged contrast-dependent SSVEP responses to each grating. When both gratings are at equal contrast the anti-phase ensures that the signals for each grating will cancel out on the scalp. When one grating has higher contrast, the 15 Hz signal will be more strongly phase-aligned to that grating, providing an index of the encoding of the sensory evidence for this experiment (i.e., the differential contrast of the two gratings). The 15 Hz phase-tagged signals were calculated by convolving the CSD data with a 15 Hz sinewave (171 samples length, 334ms, 5 cycles), and the phase was projected on to the mean phase 400ms after the initial response, when the 15 Hz signal was steady. The real component of this projection reflects the strength of the alignment with this phase, with negative values corresponding to the opposite phase, which we term the *differential SSVEP*. Mean topographies were used to pick the best channel from a cluster centred on Oz, per person. We took the mean value of this differential SSVEP from 700:1000ms after initial response. We could only analyse the continued evidence condition, as there was no post-choice visual stimulus when evidence was extinguished.

The 30 Hz SSVEP, reflecting overall stimulus engagement to both stimuli, was taken from the same Fourier transformed data described above, from the Oz electrode, and normalised to the adjacent frequencies. The mean amplitude was taken −400:-200ms before the initial response, and 700:1000ms after it.

### Data & Code Accessibility

Task scripts and anonymous data are available at https://osf.io/4dqkz/ and MATLAB code for analysing this data is available at https://doi.org/10.5281/zenodo.7550911.

## Results

### Pre-choice CPP predicts choice accuracy and simultaneously reported confidence

In experiment 1, participants made two-alternative forced choice dot motion direction discrimination decisions and simultaneously reported their confidence in that choice (Figure 1). As expected, Accuracy (β = 0.15, t (11765) = 6.75, p <.0001) and Confidence (β = 0.06, t (11765) = 7.73, p <.0001) increased with Stimulus Duration, as shown in Figure 2b-e.

**Figure 2.**
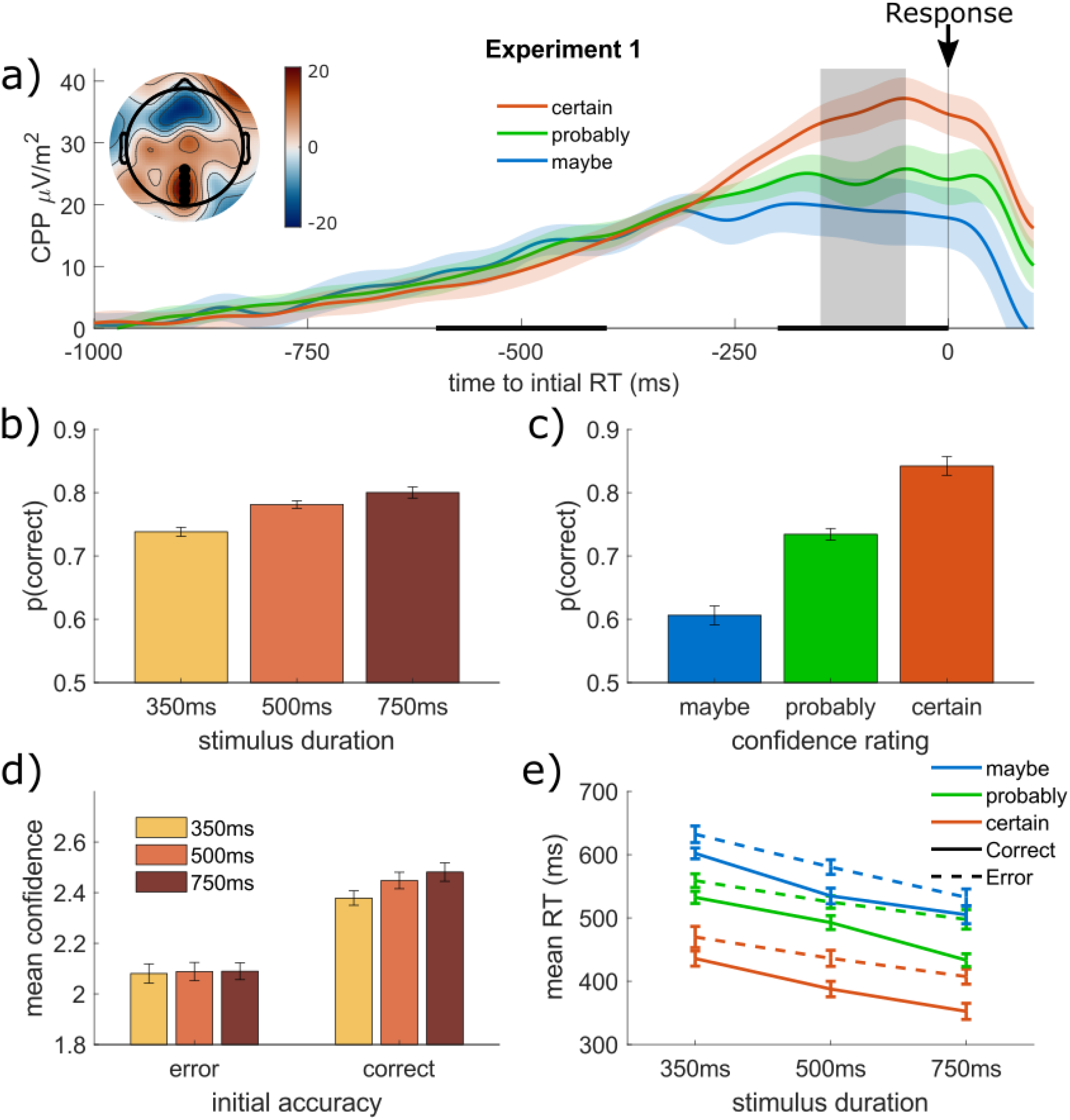
Pre-choice CPP predicts confidence reported simultaneously with direction in Experiment 1. a) Mean pre-choice CPP traces in experiment 1, leading up to response time, split by Confidence (within each stimulus duration and averaged across durations). Experiment 1 pre-choice CPP amplitude within the grey time-window is higher for ‘certain’ responses (black bar = effect of Confidence, p<.05 within each 100ms time-bin). The inset topography shows the mean activity within the time-window (red=positive, blue=negative) for the grand-mean over all confidence levels, with the black dots showing the electrodes used for CPP selection. b) Mean accuracy increases with stimulus duration. c) Mean accuracy is higher for trials rated higher confidence. d) Mean confidence rating (1 = maybe, 2 = probably, 3 = certain) increases with stimulus duration only for correct responses. e) Mean post-offset RT is quicker for longer stimulus durations, and for higher confidence ratings, but is slower for incorrect trials (dashed lines).

RT relative to stimulus offset decreased with longer Stimulus Durations (β = −0.23, t (11759) = −33.32, p <.0001), and with greater Confidence (β= −0.11, t (11759) = −15.60, p <.0001), and was faster for correct responses (β = −0.33, t (11759) = −43.47, p <.0001). There were also interactions of Accuracy and Stimulus Duration (β = −0.03, t (11759) = −4.51, p <.0001), and Accuracy and Confidence (β = −0.03, t (11759) = −4.80, p <.0001), as the effects of Stimulus Duration and Confidence on RT were stronger on correct trials.

As previously observed in studies of perceptual choice, the CPP exhibited a steady build-up during deliberation (Figure 2a), reaching a peak just before response execution. The CPP reached a significantly higher amplitude prior to Correct responses compared to erroneous ones (β = 0.04, t (11759) =3.93, p <.0001), and on trials with higher reported Confidence (β = 0.05, t (11759) =4.88, p <.0001) with no significant interactions (p > .5). Post-hoc contrasts indicated that CPP amplitude was significantly larger for ‘certain’ compared to ‘maybe’ (F (1,11761) = 8.87, p =.0029) and ‘probably’ (F (1,11761) =5.32, p =.0211) confidence ratings, but did not differ reliably between ‘maybe’ and ‘probably’ confidence ratings (F (1,11761) = 0.39, p =.53).

### Pre-choice CPP only predicts delayed confidence reports when evidence is Extinguished following commitment

In experiment 2, participants made two-alternative forced choice contrast discriminations and reported their confidence after a delay of 1000ms (Figure 1), during which the physical evidence either Continued on-screen or was Extinguished. Discrimination accuracy increased from initial-choice to final-choice (β = 0.15, t (42494) = 11.90, p <.0001, Figure 3c) with a significant Response (initial vs final) by Post-choice Evidence (Continued vs Extinguished) interaction (β = 0.05, t (42494) = 4.12, p <.0001), as accuracy increased more when evidence presentation Continued throughout the delay period.

**Figure 3.**
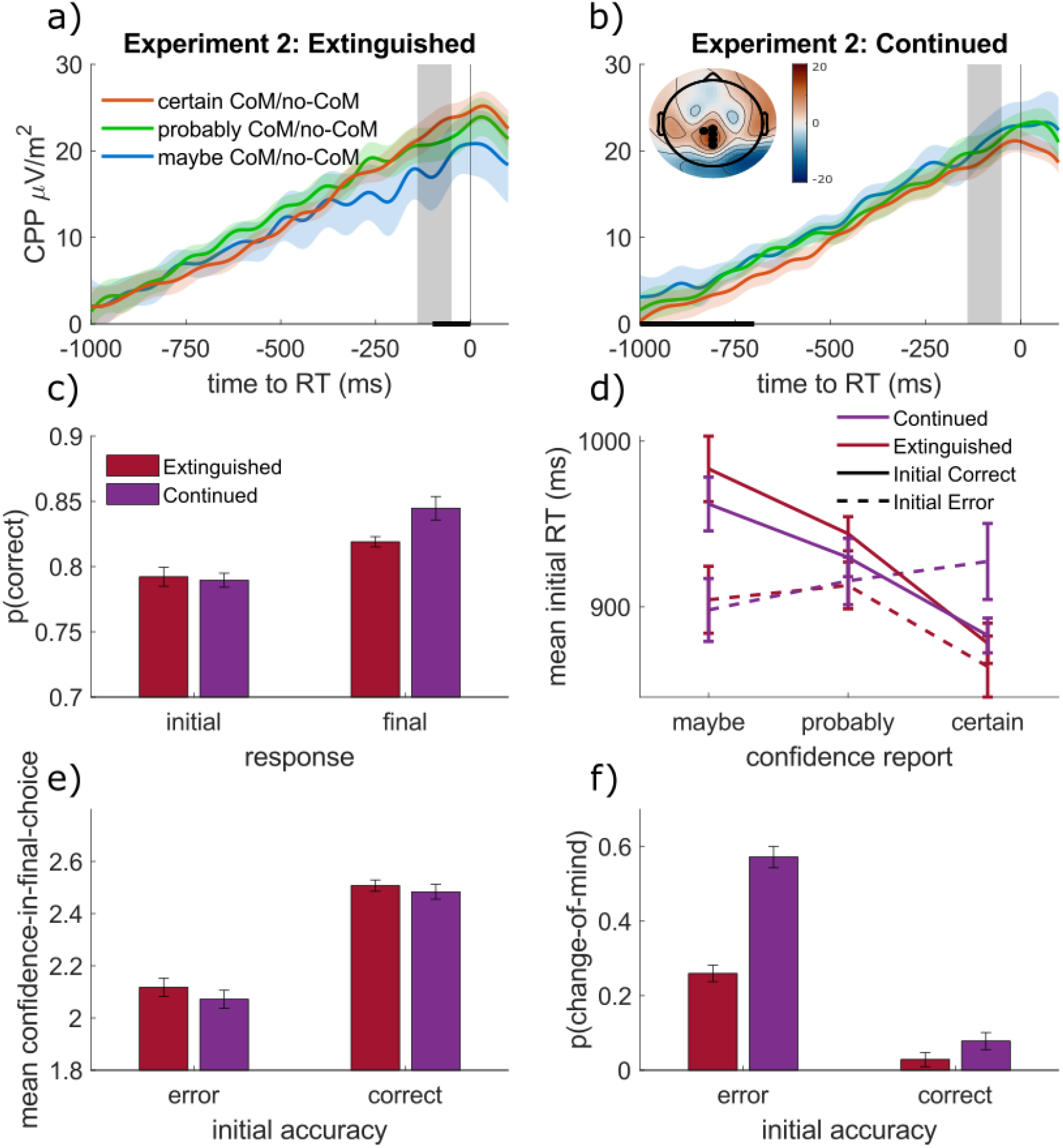
Pre-choice CPP in Experiment 2 predicts delayed confidence only if evidence is extinguished after response. To allow better comparison with Figure 2, trials with changes-of-mind are excluded from the waveforms in panels a & b only (see Supplementary Figure 2 for those trials). a) Pre-choice CPP amplitude is greater on trials later rated as higher confidence-in-final-choice, but only when the evidence is extinguished after the initial response (black bar shows significant effects of confidence on mean amplitude within 100ms time-bins, p<.05). b) If post-choice evidence instead continues, this disrupts the link between pre-choice CPP and final-choice confidence. The inset topography shows the grand-mean activity within the time-window across all trials (both evidence conditions, all confidence ratings), with the black dots showing the electrodes used for CPP selection (same scale as inset in Figure 2). c) Choice accuracy increases between initial and final responses, with a much greater increase if evidence continues. d) Mean RT for initial responses is quicker for trials rated higher confidence-in-final-choice, slower for correct responses, with an interaction of the two. e) Mean confidence-in-final-choice (1=maybe, 2=probably, 3=certain) is higher after correct responses, but slightly lower if evidence continues. f) Changes-of-mind are far more common following errors, and continued evidence increases changes-of-mind, with greater effect on error trials.

We first analysed the participants’ confidence-in-final-choice, irrespective of whether that report disagreed with the initial choice (i.e., both ‘certain left’ and ‘certain right’ responses were classed as high confidence-in-final-choice). Reported confidence-in-final-choice was higher following a Correct initial choice (Figure 3e; β = 0.22, t (21249) = 36.45, p <.0001), and lower when evidence presentation Continued throughout the delay period (β =-0.02, t (21245) =-3.86, p =.0001), with no Accuracy by Post-choice Evidence interaction (p > .1).

Investigating this trend further we found that Continued evidence led to a significant increase in change-of-mind rates (β = .54, t (16930) = 12.40, p < .0001) following correct initial response (β = .54, t (16930) = 12.40, p <. 0001) as well as erroneous ones (β = .80, t (4315) = 20.88, p <. 0001), although the effect was significantly greater for initial errors (Accuracy by Post-Choice Evidence interaction (β = −0.10, t (21245) = −4.34, p <. 0001). Thus, while post-choice evidence mainly causes corrective changes-of-mind leading to greater final accuracy overall, it also causes a reduction in confidence ratings and increase in choice reversals following correct initial responses (Figure 3c). This observation likely stems from the fact that, based on confidence levels reported for correct trials in Experiment 1 and in the extinguished evidence condition of Experiment 2, participants were close to the maximum level of confidence that could be reported at the time of initial commitment, reducing the scope for any further measurable increases.

Turning to the CPP, we first sought to test the relationship between the delayed confidence reports and its amplitude measured immediately prior to the initial choice. To allow direct comparison with the pre-choice CPP effects observed in Experiment 1 (where participants rated confidence alongside their choice and therefore could not report changes-of-mind), we excluded any trials with changes-of-mind and examined pre-choice CPP amplitude as a function of final confidence (similar effects were observed when changes-of-mind were included, see Supplementary Figure 2 for statistics). Consistent with the results reported for Experiment 1, pre-choice CPP amplitude increased with Confidence (β = 0.01, t (18512) = 2.06, p = .0394), although the effect of choice Accuracy did not reach significance (β = 0.01, t (18512) = 1.02, p = .31). There was also a Confidence*Post-choice Evidence interaction (β =-0.02, t (18512) =-2.24, p = .0251), and separate LMMs in each Post-choice Evidence condition indicated that pre-choice CPP amplitude increased with Confidence when evidence was Extinguished during the delay period (Figure 3a; β = 0.04, t (8485) = 2.74, p =.0062), but not when evidence Continued (p =.9699; Figure 3b). This accords with the observation that post-choice evidence had a substantial influence on the accuracy of the final confidence reports - if decision processes are updated during the post-choice interval, then this would naturally reduce the degree to which pre-choice CPP amplitudes would predict the confidence level reached by the end of the trial.

### Post-choice CPP scales inversely with final choice confidence irrespective of initial accuracy

Next, we looked at how the CPP evolves during the post-choice delay period in Experiment 2 (Figure 4), including in change-of-mind trials. When evidence was Extinguished the grand-average CPP returned gradually back to baseline over the next 1000ms (Supplementary Figure 1) but, when evidence presentation continued, the CPP decayed at a markedly slower rate overall, and exhibited a positive build-up during the delay period on certain trial types.

**Figure 4.**
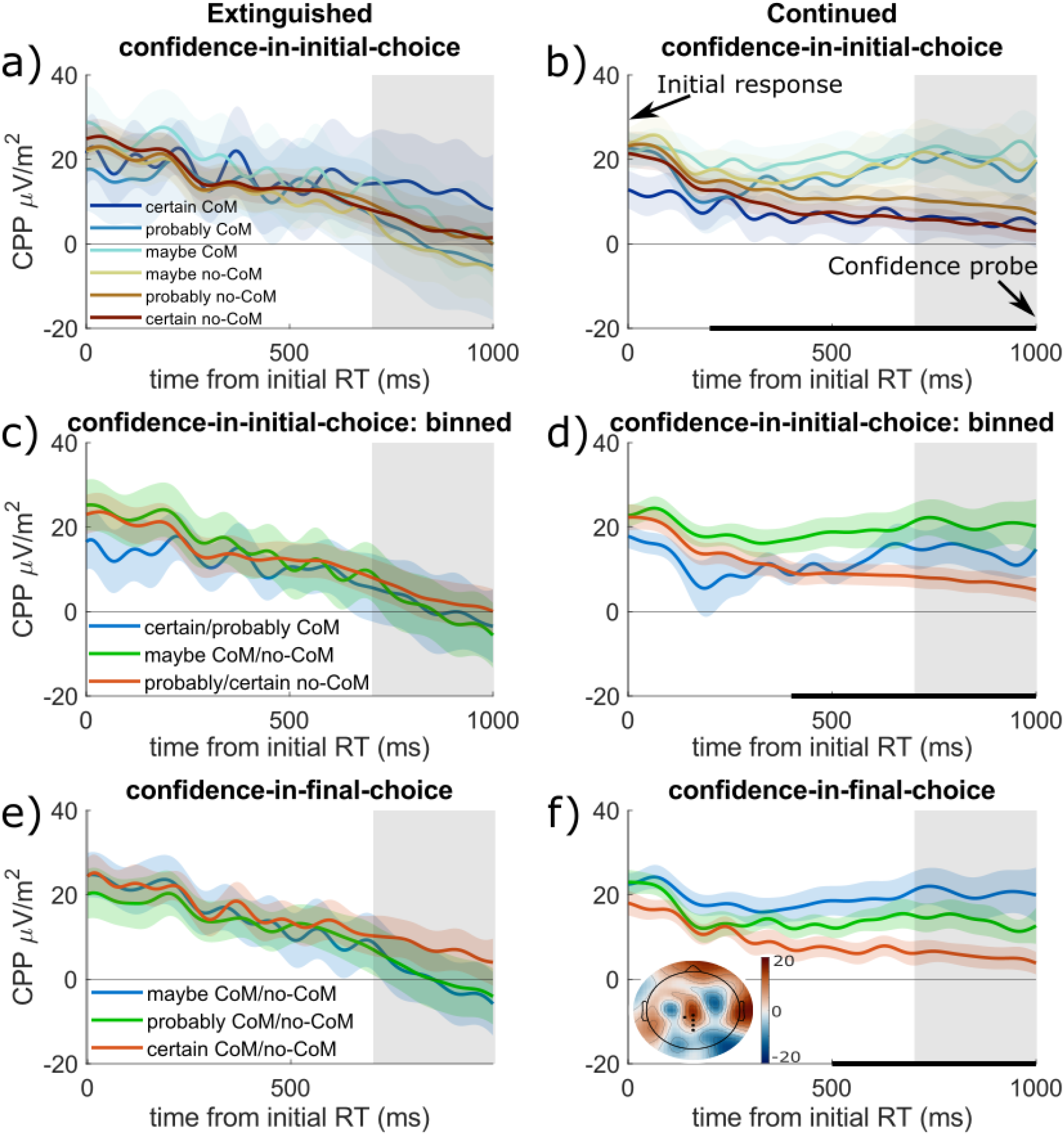
Post-choice CPP is sensitive to post-choice evidence and increases when participants change their mind or have low confidence in their final choice. Post-choice CPP refers to the CPP time-course between the initial choice response and the final response (with confidence report) 1000ms later in Experiment 2. a) The 3-point confidence scale was re-coded into a 6-point confidence scale, taking into account whether the trial included a change-of-mind (CoM) or not, indicating whether the participant believed the initial response was wrong or correct (termed confidence-in-initial-choice). The mean CPP (700:1000ms) decreased after the initial choice if the evidence was Extinguished at that time, with no significant difference depending on confidence-in-initial-choice. b) When Evidence Continued after the initial response, the post-choice CPP plateaued, and rose again on trials where subjects’ final ratings translated to their initial choice being ‘maybe’, or ‘probably CoM’. The effect of confidence-in-initial-choice became significant from about 400ms after the initial decision (solid black bars show p<.05 in 100ms time-windows). c) The same data as above, but with confidence-in-initial-choice binned in pairs to increase the trial-numbers within one waveform; again, Extinguished trials have no differences, but d) Continued trials show a significant effect, which is non-linear, as the medium bin (maybe CoM/no-CoM) is highest. e) Extinguished evidence trials did not differ by confidence-in-final-choice (i.e., maybe/probably/certain) responses. f) Continued-evidence trials had greater post-choice CPP amplitudes on trials rated ‘maybe’ than ‘probably’ or ‘certain’. The topography inset in panel f) shows the mean amplitude within the grey window for the low confidence-in-final-choice trials (red=positive, same scale as Figures 2 & 3, black dots are CPP electrodes), and the solid black bars along the bottom show which 100ms time-windows have significant effects in the CPP waveforms (p < .05, none were significant for the Extinguished conditions).

To match analyses conducted in previous studies of confidence-related modulations of post-choice ERP signals (e.g. Boldt & Yeung, 2015; Feuerriegel et al., 2022), we first analysed post-choice CPP amplitudes (measured at the end of the delay period) as a function of confidence-in-initial-choice. Here, we observed an interaction of Post-choice Evidence and Confidence-in-initial-choice (β = −0.02, t(21073) = −3.084, p = .0020), driven by higher post-choice CPP amplitudes for intermediate confidence-ratings when evidence presentation Continued during the delay period (β = −.03, t (12014) = −3.59, p = .0003), an effect that was absent when evidence was Extinguished (β = 0.00, t(9059) = 0.11, p = .91). The absence of any relationship with confidence on Extinguished trials appears to be in direct disagreement with results reported by Boldt & Yeung, (2015). However, that study used the interval immediately prior to the initial choice for its baseline correction which could cause pre-choice amplitude differences to be transferred to post-choice measurements. When we applied the same pre-choice baseline correction, we found the same pattern of results as Boldt and Yeung, with the post-choice CPP now decreasing with confidence-in-initial-choice in the Extinguished condition both in our measurement window and in the one used by Boldt & Yeung (Supplementary Figure 3). This last observation also indicates that, while the present study used confidence scales that were mapped to the choice alternatives (left vs right) and Boldt & Yeung (2015) used a scale mapped to the accuracy of the initial choice (correct vs incorrect), these differences did not alter the post-choice dynamics of the CPP qualitatively, at least in the Extinguished condition.

Although there was a significant main effect of Confidence-in-initial-choice on CPP amplitude, Figure 4b shows that CPP amplitudes were very similar for ‘certain CoM’ and ‘certain no-CoM’ trials where the difference in confidence-in-initial-choice is greatest (p=.907, BF_01_= 4.543, indicating moderate evidence for the null). CPP amplitudes were also highly similar for ‘maybe CoM’ compared to ‘maybe no-CoM’ trials (p=.890, BF_01_= 4.701, indicating moderate evidence for the null) with differences only apparent between ‘probably CoM’ and ‘probably no-CoM’ (p=.035, BF_01_= 0.573 - anecdotal evidence for the alternative hypothesis).

To investigate this pattern further, we grouped the confidence ratings such that trials were labelled according to the subjects’ confidence-in-final-choice (i.e., collapsing across left and right choices for ‘maybe’, ‘probably’, and ‘certain’). Here we observed a Confidence-in-final-choice*Post-choice Evidence interaction (β =-0.29, t (21073) = −4.22, p <.0001), driven by the fact that when evidence continued (Figure 4b), higher confidence-in-final-choice was associated with a *smaller* post-choice CPP amplitude (β =-0.02, t (12014) =-2.27, p =.0231), while no such relationship was observed when evidence was extinguished (Figure 4a; β = 0.01, t (9059) = 0.75, p =.45). BIC slightly favoured confidence-in-final-choice as a predictor of CPP amplitude over confidence-in-initial-choice (ΔBIC=-3).

Taken together, these results suggest that the post-choice CPP scales inversely with the participant’s final confidence, irrespective of whether or not the ultimately chosen alternative differs from the initially chosen one. One potential explanation for this pattern, inspired by a previously reported mathematical model (Pleskac & Busemeyer, 2011), is that the duration of post-choice evidence accumulation is confidence-dependent, such that participants are more likely to terminate the accumulation process when highly confident in a particular alternative, after which the CPP decays back to baseline as seen when evidence is Extinguished (Figure 4a). Such confidence-dependent stopping would result in lower amplitudes in our measurement window (which was toward the end of the delay period) for trials with high confidence-in-final-choice because of the earlier peak and decay of the CPP. A comparable effect was reported in Twomey et al. (2016) where the CPP was found to peak and decay earlier on trials with low difficulty when participants were required to withhold reporting dot motion direction decisions until stimulus offset (see also Rogge et al., 2022; Tagliabue et al., 2019). This pattern resulted in larger CPP amplitudes toward the end of the delay period for trials with weaker physical evidence. In other words, when CPP amplitude measurements are taken within a fixed time window within a delay period, they will scale with the proportion of trials on which evidence accumulation was still ongoing during that window (see Discussion for illustration). In the following sections we report a number of analyses designed to test the hypothesis that participants were reaching commitment earlier on trials with high final confidence.

### Earlier post-choice CPP peak latency and Confidence Reports on Trials with High Final Confidence

We ran a post-hoc exploratory analysis on the peak latency of the post-choice CPP. The surface plots in Figure 5a highlight substantial cross-trial variability in post-choice CPP peak latency and confirm that, at the single-trial level, the signal decayed rapidly after reaching its peak. Post-choice CPP peak latencies were significantly later when Post-decision Evidence Continued (Figure 5b, β=0.28, t (21245) = 4.15, p < .0001), but earlier when Confidence-in-final-choice was higher (β=-0.21, t(21245)=-2.71, p=.0067), with a significant interaction of the two (β=-0.24, t(21245)=-3.50, p=.0005). This was due to confidence only having a significant association with peak latency when evidence Continued (p=.0036), but not when it was Extinguished (p= 0.40).

**Figure 5.**
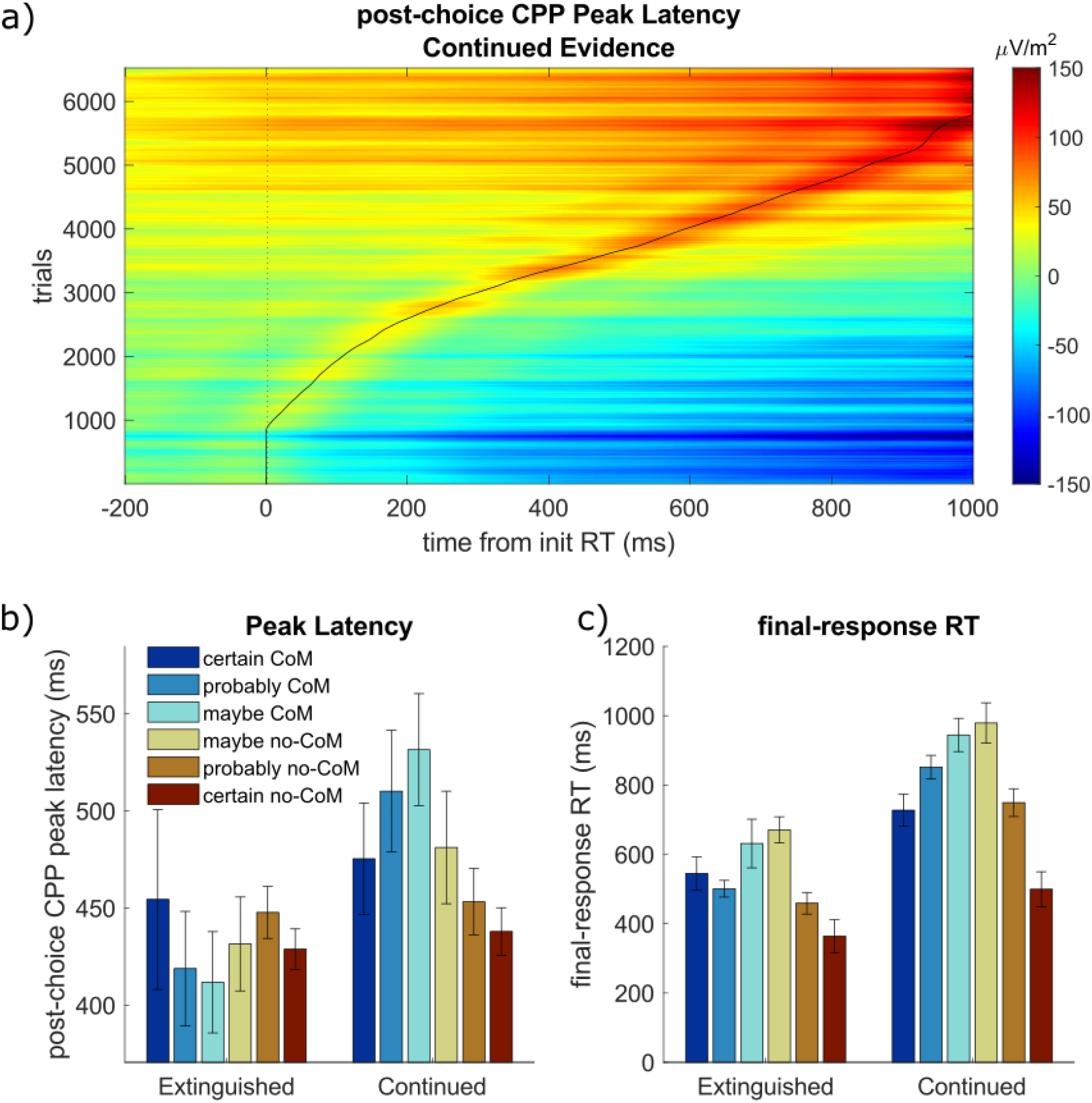
Post-choice CPP peak latency and final-response RT are quicker for high-confidence responses. a) Single-trial post-choice CPP surface plot, sorted by peak latency (curved black line) for the Continued Evidence trials (smoothed over 100 trials). The post-choice CPP decreased after the peak, rather than remaining elevated. b) The mean post-choice CPP peak latency was earlier on trials rated higher confidence-in-final-choice (darker colours), but only when evidence Continued - giving the inverted U-shaped pattern across the 6 confidence-in-initial-choice levels shown here. c) Final-response RTs showed a similar pattern when evidence Continued, with faster final-RTs for higher confidence-in-final-choice reports (darker colours). Please note that final-response RTs were cued-responses after the 1000ms delay, so did not occur around the same time as the peak latencies.

If commitment is being reached earlier on trials with high final confidence, then response times for the final confidence report should be made more quickly. Analysis of final response RTs confirmed this, with significantly faster RTs when Confidence-in-final-choice was higher (Figure 5c; β=-0.22, t(21245)=-32.44, p<.0001). RTs were also significantly faster when evidence was Extinguished (β=0.30, t(21245)=50.07, p<.0001) and there was a significant Confidence by Evidence interaction (β=-0.01, t(21245)=-2.17, p=.0298). Examining this interaction, there were significant effects of Confidence-in-final-choice both when evidence Continued or was Extinguished (p<.0001), although the effect was stronger when evidence Continued (β=-0.23 vs −0.20). As Figure 5c shows, when evidence Continued, the ‘certain CoM’ and ‘certain no-CoM’ had the fastest final-response RTs, and these were almost as fast as trials where the evidence was Extinguished (only 182ms and 135ms slower on ‘Certain CoM’ and ‘Certain No-CoM’ trials respectively, vs >290ms all the other trial types). Taken together, these results fit with the idea that deliberation stopped earlier when highly confident.

### Faster Motor Lateralisation on trials with High Final Confidence

We reasoned that, if a confidence-dependent stopping rule were being implemented, then motor preparation of the final response should emerge earlier in the delay period on trials with higher final confidence. We first examined levels of motor preparation in the same measurement window used to analyse the post-choice CPP.

There was a significant effect of Confidence-in-final-choice on mu/beta lateralisation (β=-0.07, t(21018)=-10.16, p<.0001), which significantly interacted with Post-choice Evidence (β=-0.02, t(21018)=-3.43, p=.0006). This interaction was due to a stronger effect of Confidence-in-final-choice when evidence Continued (Figure 6f; β=-0.09, p<.0001) than when Extinguished (Figure 6e; β=-0.04, p<.0001).

**Figure 6.**
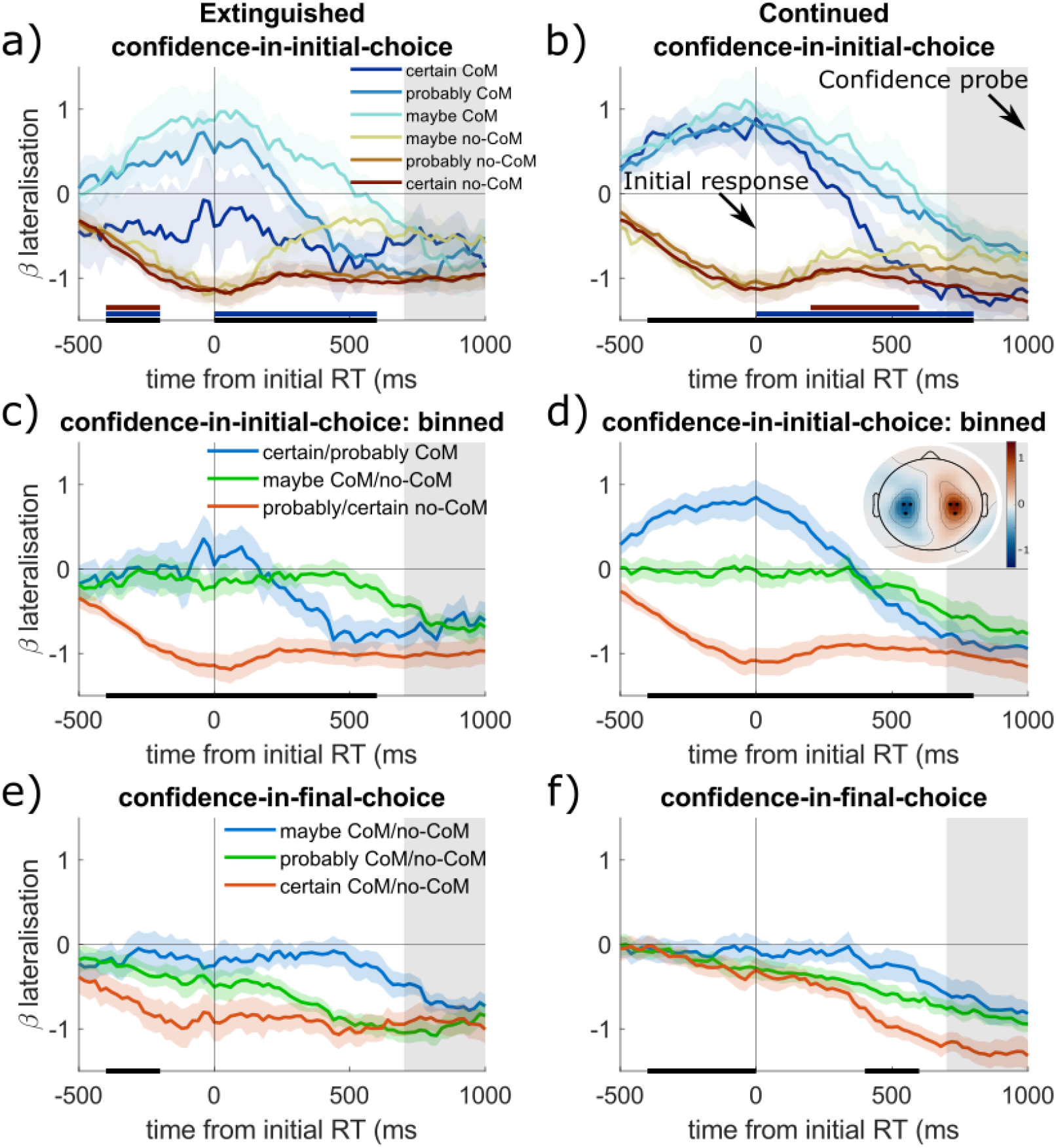
Motor lateralisation signals invert with changes-of-mind, and scale with confidence-in-final-choice. Mu/beta power lateralisation index relative to the hand used for the final response (therefore it is inverted at the time of the initial response on trials where participants changed their minds). Topography in panel d) shows the grand-mean beta lateralisation index within the grey window, averaged across all conditions and trials (red=positive, black dots = selected channels). The solid black bar shows the 200ms time-bins with significant effects (p<.05) of that factor on the **slopes** of the lateralisation, while blue/red solid bars show the same for effects of confidence-in-final-choice on **slopes** within change-of-mind/no-change in the top row. Mean mu/beta lateralisation (within the grey window, 700:1000ms after initial RT) is stronger when people have greater confidence in their initial choice, whether evidence is extinguished (a) or continued (b), and the slopes are steeper for high-confidence responses from around 400ms when evidence Continues (blue/red solid bars). When confidence-in-initial-choice is binned into adjacent pairs (c & d), it is clearer that the ‘maybe’ trials have less motor preparation, especially when Evidence Continues (d). Using this coding, the green line involves trials that do and do-not change responses, so the average is close to zero at the initial response time, however when Evidence is Extinguished (c) this is also seen in the blue line, which is due to the ‘certain CoM’ trials having slightly negative motor preparation in this time-period, perhaps reflecting motor execution errors. (e-f) Mu/beta lateralisation is stronger for trials with greater confidence-in-final-choice, and weakest for ‘maybe’ trials, especially when evidence Continued.

Motor lateralisation also varied significantly as a function of confidence-in-initial choice (β =-0.04, t (21018) =-4.90, p <.0001), although without a significant interaction of Post-choice Evidence (p > .4). Figure 6b highlights a similar but inverted pattern to that observed for the CPP, with lateralisation increasing with final confidence but not differing as a function of whether the final choice conflicted with the first. Bayesian t-tests comparing the mean amplitudes for change-of-mind and non-change with the same confidence-in-final-choice rating indicated moderate evidence for the null hypothesis for all three pairs (‘certain no-CoM’ vs ‘certain CoM’, BF_01_ = 4.451; ‘probably no-CoM’ vs ‘probably CoM’, BF_01_=3.012; ‘maybe no-CoM’ vs ‘maybe CoM’, BF_01_=4.539). Comparing the LMM including confidence-in-initial-choice to that including confidence-in-final-choice, we found that the latter again gave a better fit to the neural data (ΔBIC = −90). Thus, as predicted, greater motor preparation was observed on trials with higher reported final confidence.

To better test the hypothesis that participants were committing to and preparing their responses earlier on high final confidence trials we measured mean mu/beta lateralisation slopes in 200ms windows after the initial choice and ran separate analyses on CoM and no-CoM trials (due to the large inversion of lateralisation that occurs on CoM trials). In both CoM and no-CoM trials, trials with higher confidence-in-final-response ratings had significantly stronger slopes than lower confidence trials, and this effect was apparent from 400-800ms after initial choice (p<.05, see red & blue bars in Figure 6b). Additionally, we measured the time at which mu/beta lateralisation reached a level close to that measured immediately prior to the initial response, to see whether confidence influenced the time at which motor signals approached threshold. We took the mean mu/beta lateralisation amplitude reached for the initially chosen effector (−1 μV) and used this as a proxy for the motor commitment threshold, and then used t-tests to find when the mean waveform was no longer significantly different from this level. We did not run this analysis in the no-CoM trials as they were already at this level at the initial response time. Beta lateralisation approached this threshold value substantially earlier for ‘Certain CoM’ (480ms post-choice) compared to ‘probably CoM’ (960ms) or ‘maybe CoM’ (660ms), consistent with commitment being reached earlier on average for ‘certain’ responses. These motor lateralisation findings accord with the confidence-RT results reported above, suggesting that motor preparation during the delay period was more advanced on high-confidence trials, which would fit with a confidence-dependent stopping.

### Sensory evidence signals increase with confidence and track changes-of-mind

We also examined the post-choice dynamics of the differential SSVEP which indexes the encoding of the sensory evidence (i.e., the relative contrast of the two grating stimuli). The differential SSVEP response following the initial choice was greater on trials with higher Confidence-in-initial-choice (β=0.08, t(12102)=7.62, p<.0001), which interacted with initial Accuracy (β=0.04, t(12102)=5.85, p<.0001), as the differential SSVEP was higher following correct responses subsequently rated as ‘certain no-CoM’(β=0.11, t(9607)=8.77, p<.0001), but lower following errors subsequently rated as ‘certain no-CoM’ (Figure 7a-b; β=-0.03, t(2495)=-2.15, p=0.0313). Unlike the CPP and mu-beta signals above, confidence had a monotonic effect; as the evidence always favoured the correct response, stronger evidence encoding was associated with higher confidence in correct initial choices, and lower confidence in incorrect initial choices, i.e., greater confidence in the correct option.

**Figure 7.**
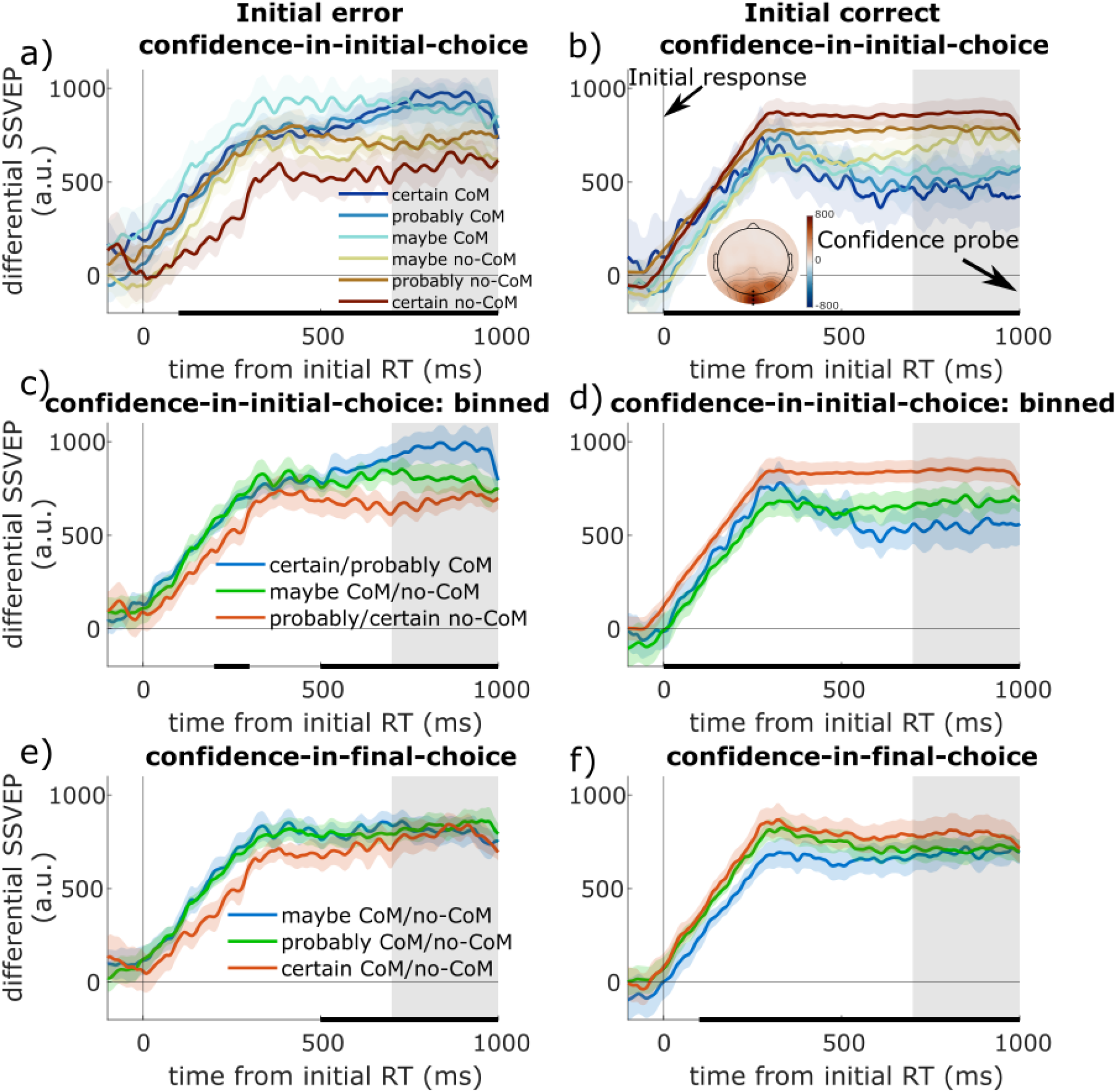
Sensory post-choice evidence signals show greater stimulus engagement with higher confidence and predict changes-of-mind. Differential SSVEP strength (arbitrary units) measures the difference in sensory signal strength for the target and non-target (i.e., the strength of sensory evidence encoding) to the post-choice continued evidence. The phase of the stimuli reset at initial RT, so the signal builds from zero then. The topography in panel b) shows the grand-mean differential SSVEP within the grey window, averaged across all Continued Evidence trials (red=positive, black dots show electrodes used for selection). Black solid bars show which 100ms time-bins have a significant effect of the factor for that initial-accuracy (p < .05). The mean differential SSVEP (within the grey time-window, 700:1000ms) shows an interaction with confidence-in-initial-choice and initial accuracy as: a) following an error response, the differential SSVEP is higher on trials where participants change their minds (blue lines), and lowest for trials in which they stick with their initial choice with high confidence (dark red); b) following a correct response, the opposite pattern is seen, with a linear increase in differential SSVEP for trials with higher confidence-in-initial-choice. c-d) When confidence-in-initial-choice is binned (to increase trial numbers for the rarer responses), the same pattern is seen. e-f) When split by confidence-in-final-choice, there was no significant interaction between initial error and confidence-in-final-choice; there was a significant positive relationship following initial correct responses, and a non-significant positive relationship following initial errors.

Confidence-in-final-choice was significantly associated with higher differential SSVEPs (β=0.06, t(12102)=7.08,p<.0001), but this had no significant interaction with initial accuracy (p=.42). BIC favoured confidence-in-initial-choice over confidence-in-final-choice as a predictor (ΔBIC=-18), in contrast to the post-choice CPP and motor preparation signals reported above.

Whereas the differential 15 Hz SSVEP traced the sensory-level representation of the relative contrast of the two gratings (i.e., the evidence), the 30 Hz SSVEP offered a metric of the overall visual response to both stimuli combined. There was no significant relationship between the post-choice 30 Hz SSVEP and Confidence-in-final-choice (p > .05) although there was a negative main effect of confidence-in-initial-choice (β=-0.03, t(11979)=-2.09,p=.0369), but this did not interact with initial accuracy (p > .05; Supplementary Figure 4) like the differential SSVEP did. BIC favoured the confidence-in-initial-choice predictor only very slightly (ΔBIC=-1). In order to interpret this lack of initial-accuracy*confidence-in-initial-choice effect in the 30 Hz signal, we directly compared it with the differential 15 Hz SSVEP. A significant three-way interaction was observed (p=.0013) indicating that the two SSVEPs differed significantly in their sensitivity to the combination of confidence and accuracy. This suggests that while the 15 Hz contrast-signal is sensitive to confidence in the correct option, the 30 Hz overall stimulus signal was not.

## Discussion

While the role of neural evidence accumulation processes in forming perceptual decisions is now well-established, the extent to which these same processes continue to operate after an initial choice to inform subsequent confidence judgments has remained an open question. The results of Experiment 1 demonstrate that when confidence is reported simultaneously with a perceptual choice, the pre-choice amplitude of a motor-independent EEG signature of evidence accumulation (CPP) increases monotonically with confidence (Figure 2a), and this effect was replicated in a different sample and task in Experiment 2 in which confidence reports were delayed by 1000ms with no intervening evidence (Figure 3a). Experiment 2 also allowed us to examine the post-choice dynamics of the CPP, and whether they were influenced by the continued availability of physical evidence.

Multiple elements of our data suggest that commitment to the final confidence report tended to be reached very soon after the initial choice in the Extinguished condition: compared to trials with continued evidence, participants had faster final RTs (Figure 5c), and no positive build-up or confidence-dependent modulation in the trial averaged post-choice CPP (Figure 4). Nevertheless, participants’ final reports were significantly more accurate than their initial choices on these trials (Figure 3c) suggesting that some further accumulation did take place, potentially drawing on evidence still in the perceptual pipeline at the time of commitment (Resulaj et al., 2009).

When the physical evidence remained on screen, the CPP did continue to exhibit confidence-dependence, but now its amplitude at the end of the delay period scaled *inversely* with the participant’s confidence in their final response (Figure 4f), irrespective of whether that entailed a change-of-mind. Previous observations led us to suspect that this trend arose from confidence-dependent variations in the duration of the post-choice CPP. Specifically, in previous studies in which participants withheld perceptual reports until the provision of a delayed response cue, the CPP has been shown to peak and decay long before response cue onset on trials with low difficulty (Rogge et al., 2022; Twomey et al., 2016). Under a standard decision model, this would occur because the boundary is reached, after which the decision variable stops accumulating and presumably decays back to baseline. As stronger evidence drives earlier decision termination, trial-averaged CPP amplitude measured late in the post-stimulus window will scale inversely with evidence strength (Rogge et al., 2022; Tagliabue et al., 2019; Twomey et al., 2016). We reasoned therefore that in the present data, post-choice accumulation may also be terminated as soon as a criterion level of confidence in one of the choice alternatives has been reached. A post-hoc exploratory analysis found that the post-choice CPP did peak significantly earlier on trials with higher confidence-in-final-choice (Figure 5b), and the peak was followed by a rapid return to baseline consistent with the process being terminated (Figure 5a). Aside from the CPP data, several other results are consistent with earlier termination of the decision process on trials with higher final confidence. First, final-response RTs were significantly faster on trials with high final-confidence (Figure 5c), consistent with actions being selected long in advance of the response cue. Second, effector-selective motor lateralisation signals exhibited earlier and greater lateralisation in favour of the ultimately chosen effector on trials with higher confidence (Figure 6b & f). The use of delayed confidence reports means we cannot tell exactly when confidence ratings were committed to, so using a free-response confidence rating may help test this prediction in the future.

While we have referred to this pattern as ‘confidence-dependent’ stopping, our data do not allow us to pinpoint the precise nature of the stopping rule that is being applied and several alternative proposals exist in the literature. According to the ‘optional stopping model’ proposed by Pleskac & Busemeyer (2011), the decision variable is translated into confidence based on its proximity to ‘confidence thresholds’ that are tied to the distinct confidence levels that can be reported, and evidence accumulation is immediately terminated when the decision variable reaches the extreme high or low confidence thresholds. This model can account for a range of behavioural effects (Moran et al., 2015) and a version of this model was recently shown to produce superior behavioural fits to a time-based stopping rule (Desender, Donner, et al., 2021). Figure 8 illustrates that the observed CPP trends can be reproduced by such a process and there are two key features of the CPP that account for this. First, the CPP is positive-going irrespective of the alternative that is favoured by the cumulative evidence (O’Connell & Kelly, 2021). Thus, the signal is expected to rise on average even when the evidence favours the alternative that was not initially endorsed. Second, the CPP decays back to baseline following choice commitment (Figures 2a, 4a, 5a), which would produce lower average amplitudes later in the delay period on trials which reached the threshold earlier (e.g. Twomey et al., 2016). Thus, the more trials which terminate early in the delay period, the more the signal decay will dominate the average signal, while the signal will build positively on average in trial types with later and/or fewer threshold crossings. Importantly, this account does not require that evidence accumulation signals evolve in the same way regardless of changes-of-mind (e.g., ‘certain CoM’ vs ‘certain no-CoM’); these trials can differ in the time taken to reach their confidence thresholds and still have decreasing *average* accumulation signals as long as the decaying signal from trials which have stopped accumulating outweigh the increasing signals from trials still accumulating.

**Figure 8.**
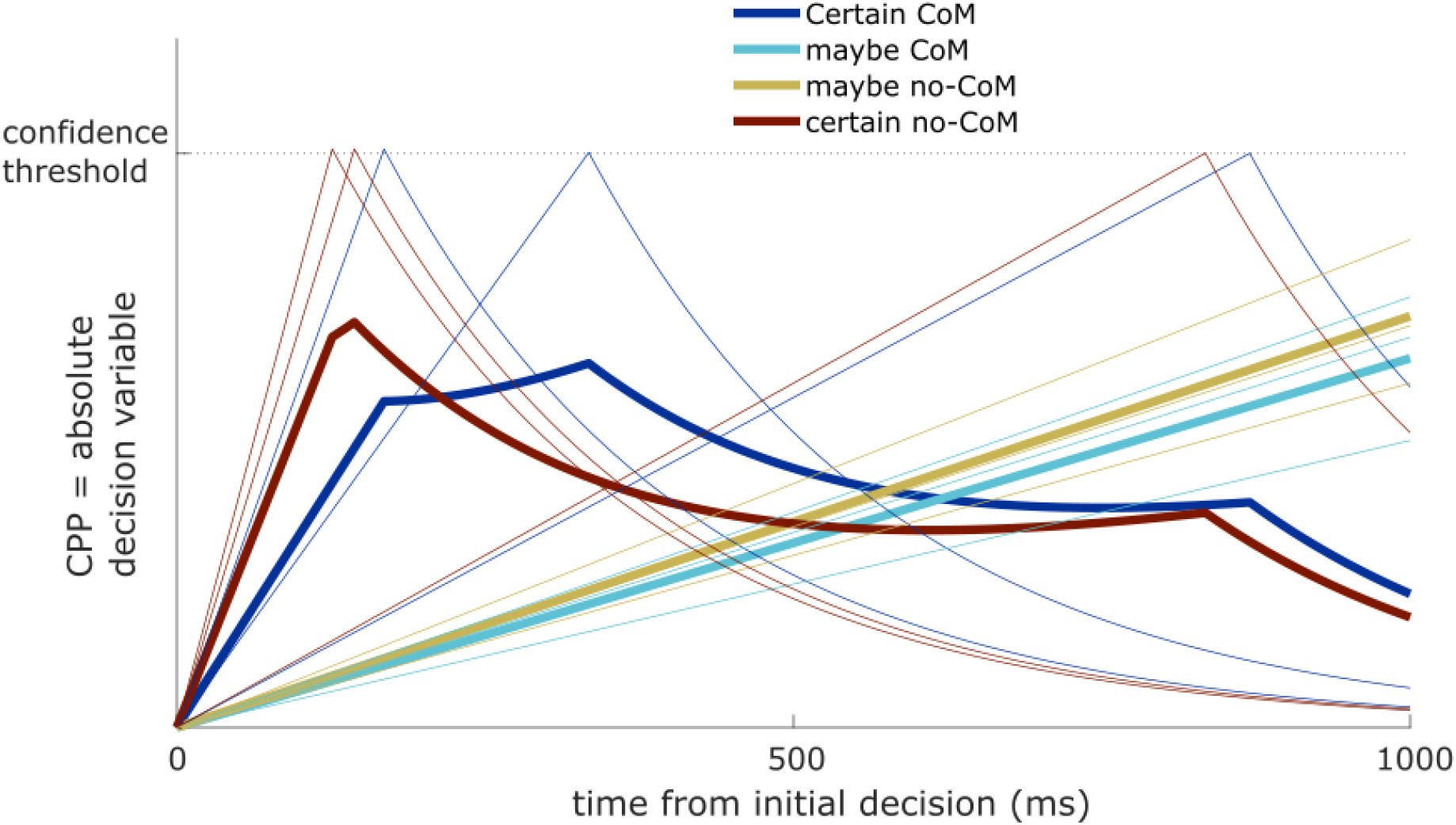
Illustration of how early confidence-boundary crossing can give a decreasing signal. Trials (thin lines) which cross the confidence-threshold (and are thus classed as ‘certain’; dark red + blue) exhibit a decay afterwards, giving a decreasing trial-average signal (thick lines). However, trials which do not reach this confidence-threshold by the end of the delay period (and thus are classed as ‘maybe’ in this simplified illustration; cyan + yellow) do not decay, thus giving a shallow positive-going trial-average signal. The CPP is represented here by the absolute decision variable, meaning that CoM and no-CoM trials both have a similar signal, although differences in the slopes (and therefore crossing-times) can still exist.

Detailed modelling, not possible given the relatively low trial numbers and fixed evidence strength in the current dataset, will be required to clarify exactly how such a stopping rule is implemented in our task. For example, in the model of Pleskac & Busemeyer (2011), while the decision variable will immediately terminate upon reaching the extreme confidence bounds, a lower probability of stopping is also assigned when passing intermediate thresholds that correspond to the intermediate confidence levels that can be reported. Alternatively, rather than terminating accumulation probabilistically upon reaching an intermediate confidence threshold, the confidence level required to terminate post-choice accumulation may decrease as a function of time (Moran et al., 2015), similar to a collapsing bound or dynamic urgency effect (Hanks et al., 2014; Yau et al., 2021). Finally, it is possible that confidence-dependent stopping could occur at the time of the initial choice, so that trials where participants were already highly confident in their initial response would have no additional accumulation at all, a possibility that would require measuring confidence at the initial choice time to investigate.

A confidence-dependent stopping rule can account for key observations in the literature on the Error Positivity (Pe), a signal that shares many functional similarities with the CPP, including its polarity and topography (Murphy et al., 2015). The Pe is typically examined in the context of tasks in which errors primarily occur due to a strong prepotency being established for one of the choice alternatives, either through differences in choice outcome probability, as in Go/No-Go tasks (e.g. Endrass et al., 2012; Niessen et al., 2017; Shalgi et al., 2009), or due to response-biases such as the antisaccade task (Falkenstein, 1990; Nieuwenhuis et al., 2001) or flanker task (Overhoff et al., 2021; Selimbeyoglu et al., 2012). It has been well-established using these kinds of tasks that the Pe is elicited by erroneous choices that are explicitly detected by participants and is greatly diminished or absent following correct choices and undetected errors (Endrass et al., 2012; Nieuwenhuis et al., 2001; O’Connell et al., 2007; Steinhauser & Yeung, 2010). These observations have fuelled the theory that the Pe reflects the operation of a process that is designed specifically to detect action errors (Desender, Ridderinkhof, et al., 2021). That is, rather than being referenced to the original choice alternatives, post-choice evidence is accumulated in a new reference frame representing the probability that the preceding choice was incorrect. However, our results highlight another plausible functional account in which the Pe can be understood as a continuation of the same process indexed by the CPP, i.e., an evidence accumulation process mapped to the choice alternatives. Taking the example of the Go/No-Go task, Go trials have much greater probability than No-Go trials and therefore the decision bounds would be expected to be much higher for the latter than the former. Consequently, for a participant to change their mind following an error of commission on a No-Go trial, post-choice evidence would need to accumulate to the higher No-Go bound, leading to a larger post-error signal.

However, our data do not exclude the possibility that the post-choice CPP reflects a metacognitive operation that is distinct from the evidence accumulation process that it traces prior to the initial choice. While we have presented the stopping-rule as a continuation of the initial decision process, it can be equally applied to a metacognitive accumulation process that evaluates the initial choice accuracy (e.g. Desender, Donner, et al., 2021; Fleming & Daw, 2016), yet stops earlier when highly confident that the response was either correct or erroneous. These two possibilities give very similar predictions for the neural and behavioural data, especially in the current paradigm and dataset, so we are not able to distinguish between them. Future experiments measuring confidence at initial and final choices, and manipulating post-choice evidence, will aim to directly compare these two theories.

To our knowledge, our study is the first to examine the post-choice CPP in the presence of additional ambiguous evidence, but two previous studies have examined it on tasks in which a delay was imposed between initial choice and confidence reporting with no intervening evidence (Boldt & Yeung, 2015; Feuerriegel et al., 2022). In the case of Boldt & Yeung (2015), the post-choice parietal ERP was found to scale negatively and monotonically with confidence-in-the-initial choice, contrary to the present results. However, in that study, signals were baseline-corrected to an interval immediately prior to the initial choice, which potentially confounds pre- and post-choice variations in cumulative evidence. When Feuerriegel et al. (2022) applied a pre-stimulus baseline correction, these post-commitment confidence effects on amplitude were substantially diminished and observed only following erroneous choices. We found a similar pattern here, with the application of a pre-response baseline period producing significantly greater post-choice CPP amplitudes for low confidence-in-initial-choice ratings, which were not observed with a pre-stimulus baseline (see Supplementary Information).

We also found that the encoding of the sensory evidence (differential contrast) in early visual areas, as indexed by the differential SSVEP, was highly sensitive to confidence (Figure 7a & b). Unlike the CPP and motor lateralisation signals, the differential SSVEP scaled most strongly with the participant’s confidence relative to the correct option, with stronger responses on initially correct trials that were rated as likely to be correct and weaker responses on those rated as likely to be incorrect (the opposite pattern was seen following initial errors). Previous studies have found sensory ERPs are stronger on higher confidence trials (Squires et al., 1973; Zakrzewski et al., 2019), but to our knowledge this is the first finding of the SSVEP showing a similar scaling. The differential SSVEP signals remained at a relatively constant level throughout the post-choice delay period even on trials with high final confidence suggesting that early termination of the decision process did not result in a disengagement of visual processing resources from the stimulus. It was not possible in the present study to establish whether these signal modulations by confidence manifest before or after commitment to the final confidence report. Again, the use of speeded confidence reports would be useful in this regard.

In conclusion, our data suggest that evidence accumulation persists following initial choice commitment when physical evidence remains available and is terminated in a confidence-dependent manner. Whether post-choice evidence accumulation is mapped to the choice alternatives or is reframed to evaluate the accuracy of the initial choice remains an open question. In general, our study highlights an important methodological consideration when measuring post-commitment CPP activity; trial-averaged amplitudes can vary as a function of the duration of the accumulation process as well as the bounds associated with each of the choice alternatives. Using paradigms in which participants make speeded, rather than delayed, post-choice confidence reports may facilitate the acquisition of CPP measurements that are more reflective of the state of the decision process at the time the participant commits to their final confidence report.

## Acknowledgements

J.G and R.G.O. were supported by Horizon 2020 European Research Council Consolidator Grant IndDecision 865474. S.P.K. was supported by Science Foundation Ireland (15/CDA/3591) and The Wellcome Trust (219572/Z/19/Z)

## Notes

Conflicts of interest: The authors declare no competing financial of interest.

https://osf.io/4dqkz/

https://doi.org/10.5281/zenodo.7550911

### Competing Interest Statement

The authors have declared no competing interest.

