## Supplementary Materials for "Confidence is predicted by pre- and post-choice decision signal dynamics"

#### Effect of Continued Evidence on Post-choice CPP

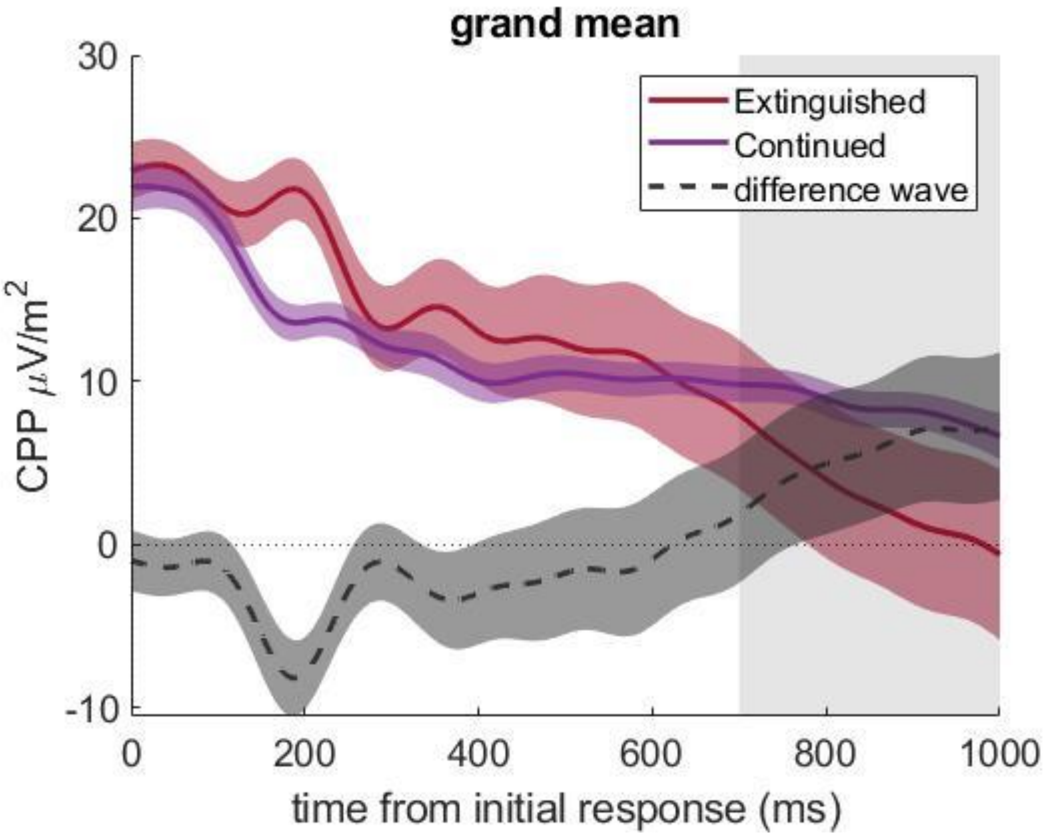


***Supplementary Figure 1. Post-choice CPP when evidence Continues or is Extinguished after the initial choice in Experiment 2.*** *The mean post-choice CPP waveform builds-up prior to the initial response, before returning towards baseline over the 1000ms delay. The Continued and Extinguished conditions differ around 190ms after initial choice, likely due to a visual potential from the stimulus-offset in the Extinguished condition, and again from 700ms onwards as the Continued condition has a much shallower decrease in amplitude over the delay.*

#### Including change-of-mind trials from confidence analyses

In the main text, for experiment 2 we excluded change-of-mind trials from the confidence analysis of the pre-choice CPP, to more closely match experiment 1 where people could not change their minds (and their confidence reports). Here we present the Experiment 2 pre-choice CPP with changes-of-mind included.

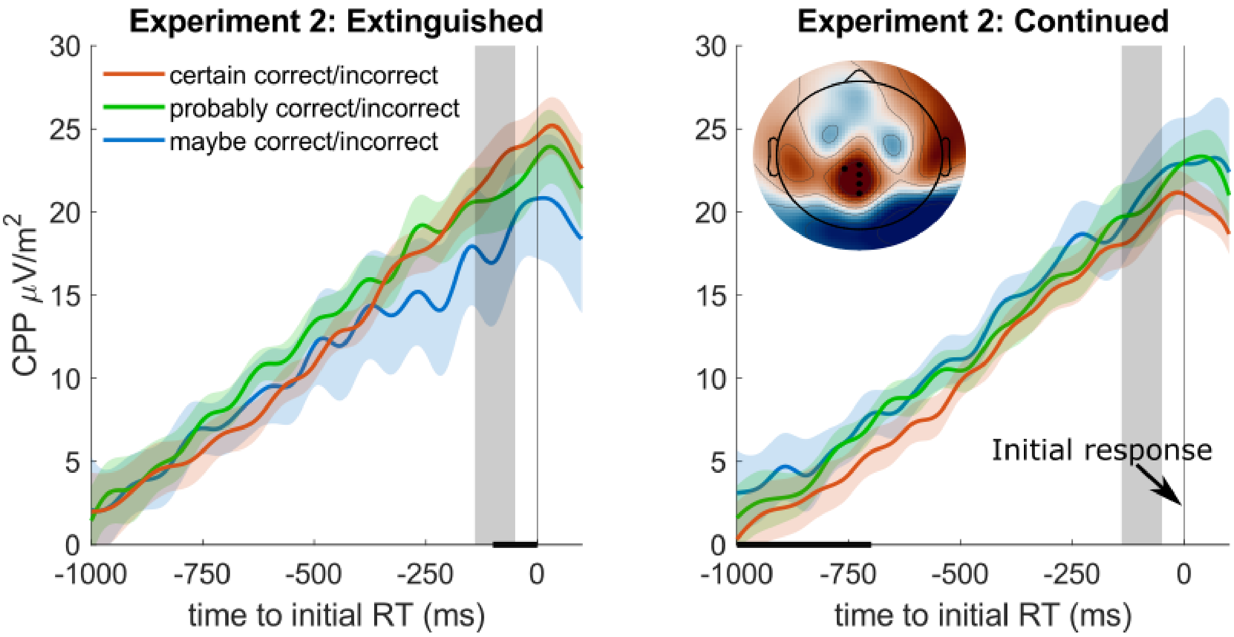


***Supplementary Figure 2. Pre-choice CPP in Experiment 2, with change-of-mind trials not excluded.*** *a) When evidence is extinguished, there is still a significantly higher pre-choice CPP amplitude for 'certain' responses, which is not seen when evidence continues (b). The topography shows the grand-mean CSD in the grey time-window (collapsing over both evidence conditions and all confidence ratings).*

The pre-choice CPP was not affected by Confidence if Change-of-mind trials were included (β = 0.01, t (21245) = 1.35, p =.18), but there was still a Confidence*Evidence interaction (β =-0.01, t (21245) =-2.19, p =.0286). Separate LMM in each Evidence condition found that when Evidence was extinguished, Confidence increased with pre-choice CPP (Figure S6a; β = 0.02, t (9141) = 2.30, p =.0214), while there was a non-significant negative relationship when evidence continued (Figure S6b; β =-0.00, t (12104) =-0.41, p =.68).

### Pre-response baseline-correction on the Post-choice CPP

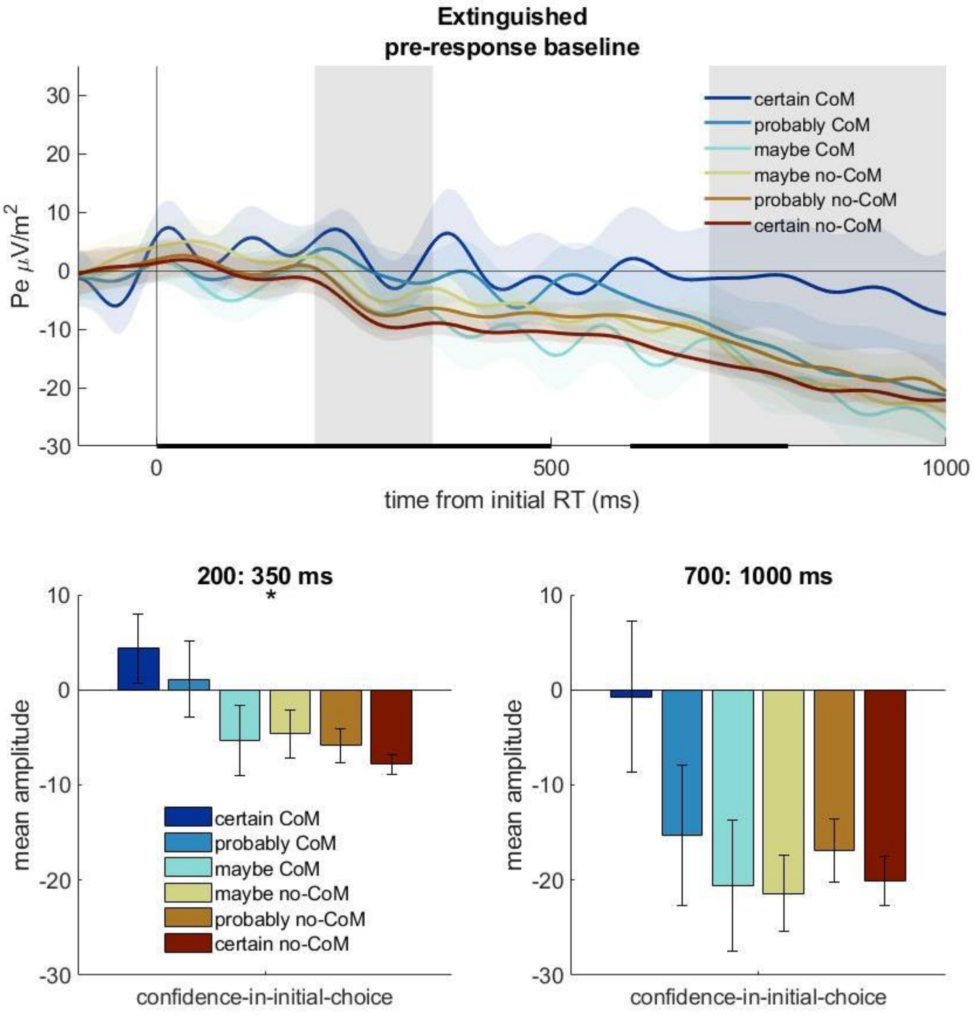


***Supplementary Figure 3. Post-choice CPP using a pre-response baseline.*** *The top panel shows the pre-response (100:0ms) baseline-corrected waveform (Extinguished Post-choice Evidence only) during the delay period. The black bars at the bottom of the waveforms show 100ms bins with significant effects of confidence-in-initial-choice (p<.05), and the stars in the bottom panels reflect significant effects within that post-choice evidence condition (p<.05). a) Using a pre-response baseline, the post-choice CPP does not rise much after the initial response but decreases more sharply when confidence-in-initial-choice was higher. b) The mean amplitudes in the early window (200:350ms) baseline show a decrease in amplitude as confidence-in-initial-choice increases (p<.05). c) There is not a significant relationship within the later time-window (700:1000ms), although the effect is still in the same direction (p=.095).*

### 30 Hz SSVEP figures

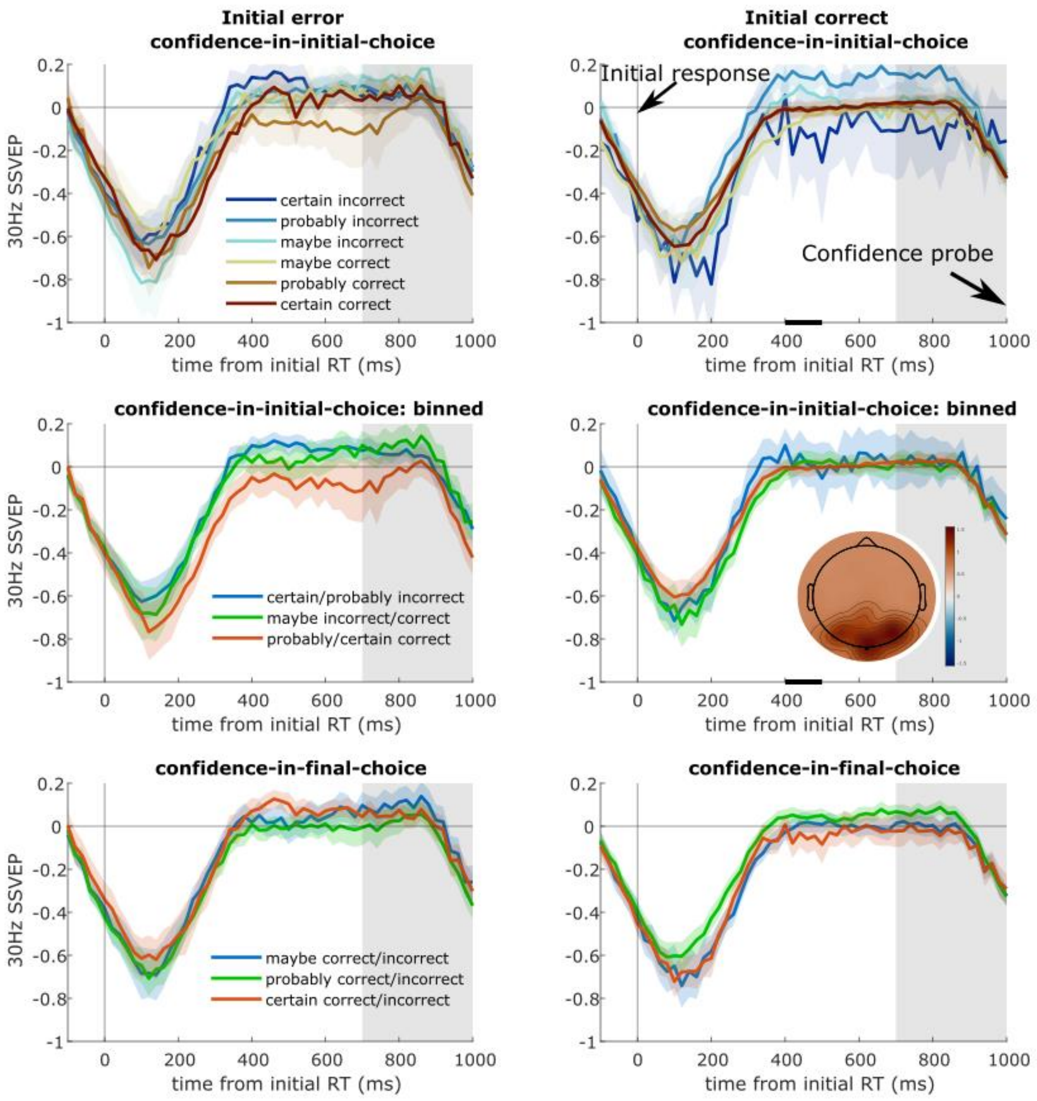


***Supplementary Figure 4. Post-choice stimulus encoding SSVEP.*** *The 30 Hz SSVEP power measures sensory encoding of the entire visual stimulus, so overall for both stimuli. It dips around the time of the initial response as the stimulus is reset at this time, so the SSVEP resets to be time-locked to the response. It rises back to the baseline (-500:0ms before evidence onset - the baseline stimulus was already on-screen for 300ms before this, so had reached asymptote before the contrast change), which reflects a return to pre-evidence onset strength, and decreases towards the end as the windows include time after 1000ms when the evidence is replaced by the confidence cue. The black bars show significant effects of the confidence factor in 100ms time-windows, for Continued Evidence trials following errors or correct initial responses separately (p<.05). There were no significant effects within the grey window-of-interest. The topography shows the mean SSVEP (not baselined) across all Continued Evidence trials within the window-of-interest. a) Trials split by confidence-in-initial-choice following an initial error response, and b) following initially correct responses. c-d) Trials for different confidence-in-initial-choices binned. e-f) Confidence-in-final-choice also had no significant effect.*

#### Effect of Initial Response Accuracy on Post-choice CPP

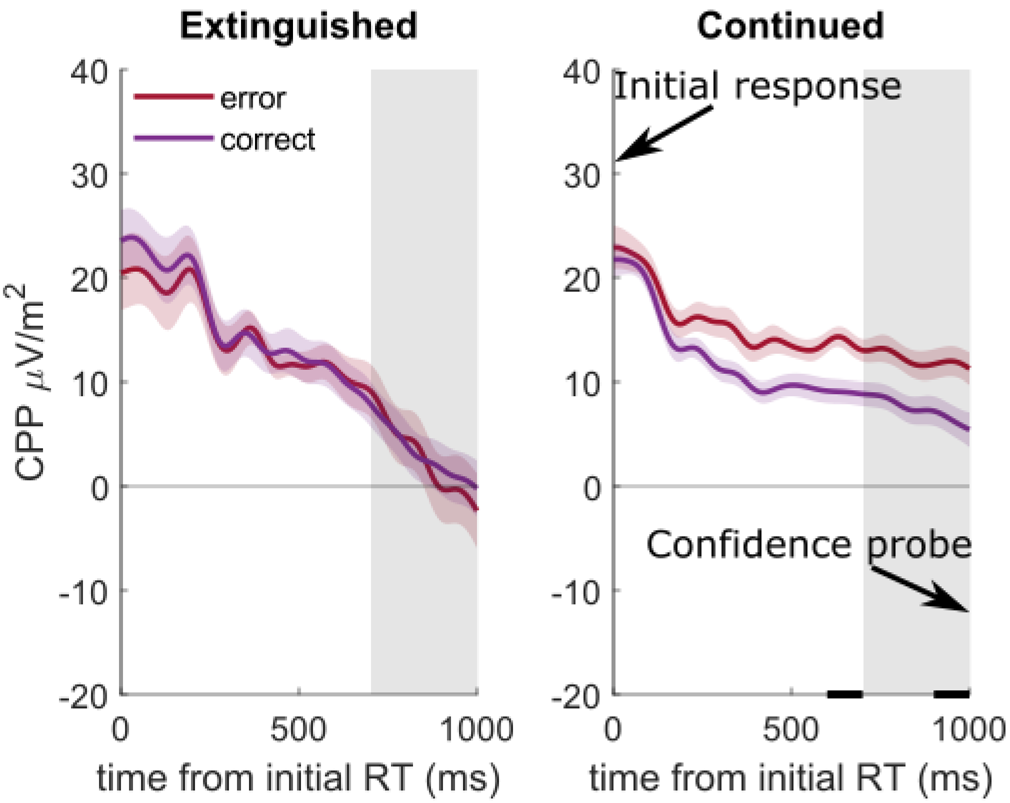


***Supplementary Figure 5. Post-choice CPP is not significantly affected by initial choice accuracy.*** *a) The mean CPP decreases after the pre-choice decision if the evidence is extinguished then, with no difference between initially correct or initially incorrect trials. b) If evidence continues, there is less of a decrease, especially following errors. LME on the mean amplitude within the grey window found a significant interaction of initial accuracy with post-choice Evidence (p = .0303), but the BIC for this model favoured both confidence-in-initial-choice and confidence-in-final-choice over this factor), although there were two 100ms time-windows with significant differences when evidence continued (solid black bar, p<.05). Given initial errors were more likely to have lower confidence and changes-of-minds (see Figure 3), the difference in the waveforms here is likely due to those factors, given the better explanatory power they have.*
